# An Aurora kinase A-BOD1L1-PP2A B56 Axis promotes chromosome segregation fidelity

**DOI:** 10.1101/2023.08.06.552174

**Authors:** Thomas J. Kucharski, Irma M. Vlasac, Martin R. Higgs, Brock C. Christensen, Susanne Bechstedt, Duane A. Compton

## Abstract

Cancer cells are often aneuploid and frequently display elevated rates of chromosome mis-segregation in a phenomenon called chromosomal instability (CIN). CIN is commonly caused by hyperstable kinetochore-microtubule (K-MT) attachments that reduces the efficiency of correction of erroneous K-MT attachments. We recently showed that UMK57, a chemical agonist of MCAK (alias KIF2C) improves chromosome segregation fidelity in CIN cancer cells although cells rapidly develop adaptive resistance. To determine the mechanism of resistance we performed unbiased proteomic screens which revealed increased phosphorylation in cells adapted to UMK57 at two Aurora kinase A phosphoacceptor sites on BOD1L1 (alias FAM44A). BOD1L1 depletion or Aurora kinase A inhibition eliminated resistance to UMK57 in CIN cancer cells. BOD1L1 localizes to spindles/kinetochores during mitosis, interacts with the PP2A phosphatase, and regulates phosphorylation levels of kinetochore proteins, chromosome alignment, mitotic progression and fidelity. Moreover, the *BOD1L1* gene is mutated in a subset of human cancers, and *BOD1L1* depletion reduces cell growth in combination with clinically relevant doses of taxol or Aurora kinase A inhibitor. Thus, an Aurora kinase A-BOD1L1-PP2A axis promotes faithful chromosome segregation during mitosis. (175 words)

## Introduction

Faithful chromosome segregation during cell division relies on proper bioriented attachment of kinetochores to spindle microtubules. Errors in kinetochore-microtubule (k-MT) attachment arise frequently during early phases of mitosis and the persistence of these errors is a common cause of the elevated rate of chromosome mis-segregation resulting in chromosomal instability (CIN) and aneuploidy in solid tumors ^1–5^. We previously demonstrated that strategies that destabilize k-MT attachments promote faithful chromosome segregation in CIN cancer cells by increasing the correction rate of k-MT attachment errors. Specifically, overexpression of either MCAK (alias KIFC) or KIF2B microtubule depolymerases suppressed chromosome mis-segregation in a variety of CIN cancer cell lines^5–8^. These data demonstrate the potential of suppressing CIN as a strategy for cancer treatment.

To pursue this strategy we recently characterized a cell-permeable small molecule agonist of MCAK activity called UMK57. MCAK plays a key role at centromeres/kinetochores by driving microtubule detachment to increase k-MT turnover to facilitate error correction prior to anaphase^9,10^. At moderate dosage (100 nM), UMK57 increases the rate of k-MT detachment and therefore decreases the rate of chromosome mis-segregation in anaphase. Surprisingly, high rates of chromosome mis-segregation returned to CIN cancer cells after only ∼72 hours of treatment with UMK57. This resistance arose within the entire cell population, did not depend on the activity of drug efflux pumps, did not occur due to a loss of drug activity, changes in the level or localization of MCAK, or changes to overall spindle morphology, and data demonstrated that MCAK activity continued to be activated by UMK57 in treated cells. Thus, this represents adaptive resistance (as opposed to acquired resistance which relies on the emergence of rare genetic variants) to UMK57 whereby changes to cell signaling pathways bypass increased MCAK activity.

To determine the mechanisms by which CIN cancer cells develop resistance to UMK57, we performed unbiased proteomic screens using mass spectrometry to identify changes in total protein and protein phosphorylation levels between acutely treated and adapted cells in mitosis. These screens revealed changes in phosphorylation of two Aurora kinase A sites in the C-terminus of the large nuclear protein BOD1L1. Subsequent studies revealed that both Aurora kinase A activity and BOD1L1 expression are required for cellular resistance to prolonged UMK57 treatment. Moreover, BOD1L1 localizes to the mitotic spindle and kinetochores, interacts with the Protein Phosphatase 2 (PP2A) complex and is required for setting correct levels of kinetochore phosphorylation to support faithful chromosome segregation. Therefore, cells rely on an Aurora kinase A-BOD1L1-PP2A axis to maintain proper k-MT attachment stability for faithful chromosome segregation and utilize this axis to alter kinetochore phosphorylation as a mechanism to adapt to persistent k-MT destabilization. Importantly, BOD1L1 mutation is associated with improved prognosis in uterine endometrial cancer. Accordingly, we show that loss of BOD1L1 expression reduces cellular fitness of cancer cell lines in culture and induces catastrophic levels of chromosome mis-segregation in combination with taxol and Aurora kinase A inhibitor.

## Results

### Proteomic analysis of UMK57 resistance in mitotic cells

To investigate the cellular mechanisms promoting resistance to UMK57, we developed conditions for proteomic screens in mitotic cells (Figure 1A). We utilized SW620 (lymph node metastasis of colon cancer) cells for this purpose because they are CIN and they undergo adaptive resistance to UMK57^11^. Specifically, SW620 cells naïve to UMK57 display lagging chromosomes in 34% of anaphases, on average. The rate of lagging chromosomes in anaphase decreases to 25% upon treatment with 100nM UMK57 for 1 hour, and rebounds to 37% following 72 hours continuous treatment (Figure 1B). SW620 cells also display reduced phosphorylation of MCAK pS95 and Aurora kinase B pT232 following 72 hours continuous treatment with UMK57 consistent with our previous findings indicating that changes in centromere/kinetochore phosphorylation levels correlated with the adaptive resistance response (Figure S1A-D). Therefore, SW620 cells likely adapt to UMK57 by altering signaling networks leading to changes in the phosphorylation of specific kinetochore proteins, as suggested previously^11^.

**Figure 1:**
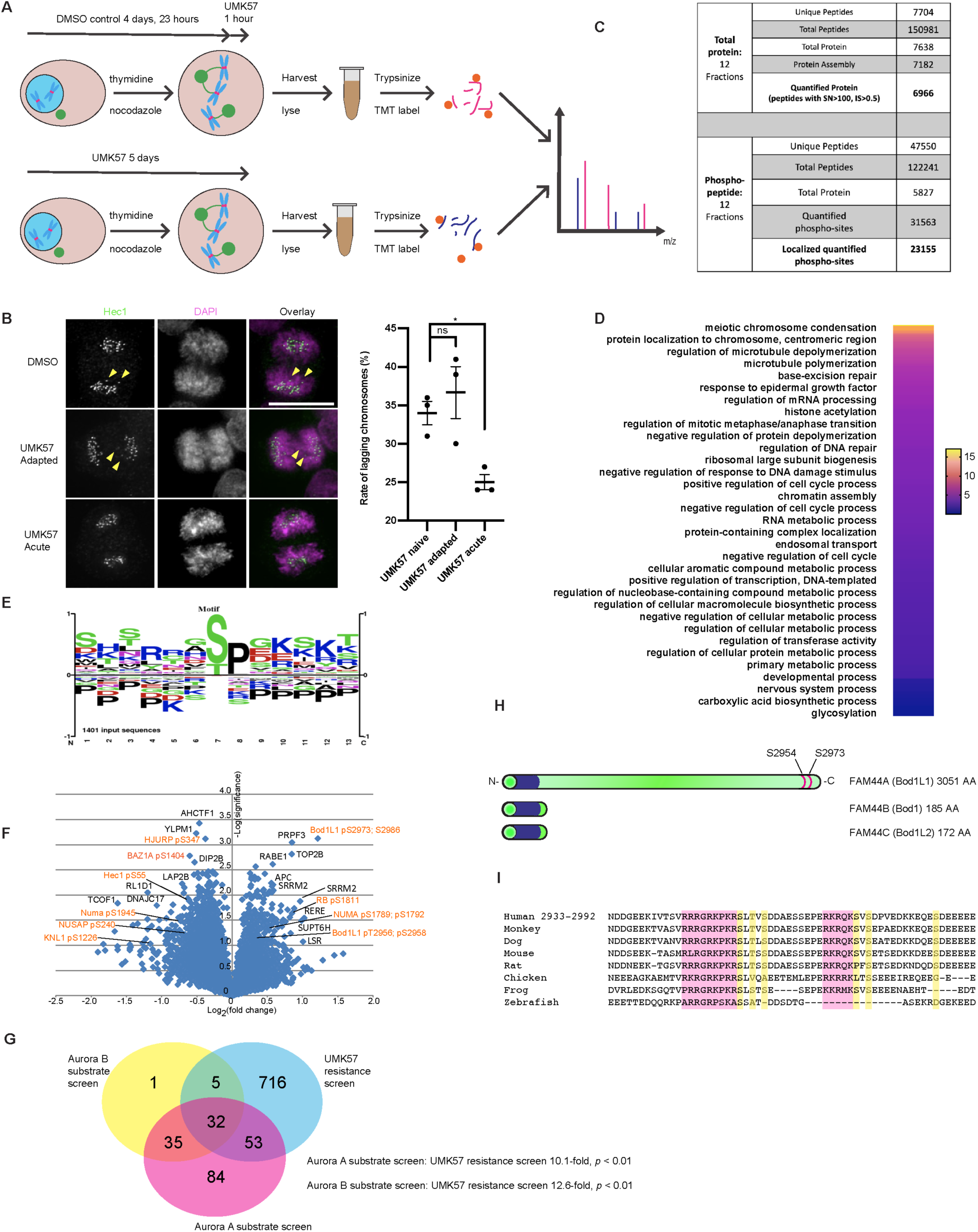
Proteomic screening reveals BOD1L1 as a mediator of UMK57 adaptation. A) Cartoon illustration depicting the experimental scheme used to discover mediators of UMK57 adaptation. SW620 cells were first treated for 3 days with UMK57 or DMSO control. They were then synchronized by thymidine-nocodazole treatment. The DMSO control sample was then treated with UMK57 for the final hour before harvest. The cells were then collected and analyzed by mass spectrometry. B) SW620 cells were prepared as in (A) but re-plated on glass coverslips instead of synchronization. After 24 hours, the cells were fixed and stained for Hec1 and DAPI. The rate of lagging chromosomes was then assessed. Statistical significance was calculated using Dunnett’s multiple comparison test. Representative images from 3 independent experiments are shown. The images were adjusted evenly for brightness and contrast for presentation. The scale bar is 10μm. C) SW620 cells were prepared as in (A) and analyzed by mass spectrometry. The number of peptides and proteins detected in the screens are shown. D) Proteins from the phospho-proteomic screen that were significantly changed (*p* < 0.1) by Student’s *t*-test were analyzed for gene ontology. The data were then transferred to Prism software and plotted. E) Proteins from the phospho-proteomic screen that were significantly changed (*p* < 0.1) by Student’s *t*-test were analyzed by PhosphositePlus for sequence motif enrichment and plotted. F) Volcano plot analysis of the phospho-proteomic screen results. Proteins of potential interest are highlighted in orange. Statistical significance was calculated using two-tailed *t*-tests. G) Analysis of overlap between the phospho-proteomic study presented here and a previous study^14^. Proteins with changes in phosphorylation greater than 4-fold caused by Aurora A or Aurora B kinase inhibitors were considered targets of Aurora A or Aurora B kinase. The list of Aurora A and Aurora B targets was compared to the list of proteins from the phospho-proteomic screen that had a significantly changed phosphorylation site (*p* < 0.1) by Student’s *t*-test. H) Cartoon drawing of the FAM44 protein family, showing the Bod1 similar domain and potential Aurora A kinase sites on BOD1L1. Scales are approximate. I) Amino acid sequence alignment of BOD1L1 orthologs from residues 2933-2992 (relative to the human sequence). Residues highlighted in pink indicate Aurora kinase defining motifs. Residues highlighted in yellow indicate potential phosphorylation sites.

For proteomic analysis, we first treated SW620 cells with 100nM UMK57 for 72 hours to represent the adapted condition. A parallel set of cells was treated with DMSO for 72 hours. The cells were then synchronized by thymidine-nocodazole (still in the presence of DMSO/UMK57) followed by 1 hour treatment of the DMSO population with 100nM UMK57 immediately prior to harvest to represent the acutely treated condition. Mitotic cells were harvested by shake-off and compared for differences in total protein level and protein phosphorylation level by mass spectrometry (Figure 1A).

For comparison of total protein levels, we obtained data for 6966 proteins based on 7704 unique and 150981 total peptides (Figure 1C). Among these, protein levels between the two cell populations varied from +1.725 to 0.41-fold change with 6813 proteins (97.7%) varying between the two cell populations within +1.2 and 0.8-fold-change. Protein levels skewed towards increased levels in one biological replicate resulting in a skewed overall distribution (Figure S1E, supplemental table 1). Among the proteins identified, the levels of Sgo2 were significantly increased (#38, +1.30-fold, *p* = 0.0037, 6 peptides) in adapted cells. Sgo2 is an interesting candidate for involvement in UMK57 adaptation as it is extensively implicated in setting kinetochore phosphorylation levels^12,13^. However, independent validation of a change in Sgo2 protein level by immunoblot was inconclusive (Figure S2F). The catalytic subunit of PP2A (PPP2CB) appeared to be decreased in protein level in adapted cells, although that was based on a single peptide and so was discarded (supplemental table 1). Thus, total protein levels in mitotic SW620 cells that have adapted to UMK57 change very little, and generally within a narrow range, relative to mitotic SW620 cells that are acutely treated with UMK57.

We next examined the phospho-proteomic changes in mitotic SW620 cells that have adapted to UMK57 versus cells that are acutely treated. We obtained data from 47,550 unique peptides from 122241 total peptides. Of these, 31,563 sites were quantified, and 23,155 sites were both quantified and localized (Figure 1C, supplemental table 2). Among these, phospho-peptide levels between the two cell populations varied from 2.98 to 0.27-fold change with 19973 (86.2%) varying between the two cell populations within 1.2 to 0.8-fold-change. 6378 phospho-peptides displayed an increased abundance in UMK57 adapted cells and 16777 peptides showed decreased abundance.

We performed Gene ontology analysis on the group of proteins that displayed significant changes in peptide phosphorylation (*p* < 0.1). We included both proteins showing increases or decreases in protein phosphorylation to capture changes caused by altered kinase or phosphatase activity. This analysis revealed significant over-representation of proteins involved in mitotic processes including chromosome condensation, centromere/chromosome localization, and microtubule polymerization and de-polymerization, consistent with an adaptive resistance to UMK57 altering mitotic processes including microtubule dynamics (Figure 1D). STRING protein-protein interaction network analysis of the top and bottom 100 proteins displaying the largest, significant changes in peptide phosphorylation revealed a cluster of proteins involved in mitosis. Of these, proteins with peptides displaying significantly increased levels of phosphorylation in adapted cells relative to acutely treated cells include Numa, 53BP1, Ska3, INCENP and BOD1L1 (Alias FAM44A). Proteins with peptides displaying significantly decreased levels of phosphorylation in adapted cells relative to acutely treated cells included Knl1, Numa, Hec1, NUSAP, INCENP and TPX2 (Figure 1F, supplemental table 2). Thus, our dataset contains proteins whose phosphoregulation might plausibly mediate adaptation to UMK57.

Analysis of the amino acid sequences of the potential phosphoacceptor sites identified in the screen with significant changes in phosphorylation level (*p*<0.1) shows over-representation of the canonical Cdk phosphorylation site motif (S/TPxxK) and possibly Aurora kinase motifs (R/KR/KxS/T) (Figure 1E). We explored if there was enrichment of Aurora substrates in our dataset utilizing a previous screen that identified many Aurora-dependent phosphoacceptors sites in mitosis^14^ and that categorized peptides as being selectively phosphorylated by either Aurora kinase A or Aurora kinase B using selective chemical inhibitors of those kinases. This analysis revealed a 10.1- and 12.6-fold over-representation of Aurora kinase A substrates and Aurora kinase B substrates, respectively, among the population of phosphoacceptors sites identified in our dataset, although many phosphoacceptors sites could be phosphorylated by both kinases (Figure 1G).

Among all proteins that displayed changes in phosphorylation level in adapted cells relative to acutely treated cells, BOD1L1 demonstrated the most significant change. 32 different BOD1L1 phospho-peptides were detected in the phospho-proteomic screen, of which 3 showed statistically significant positive changes, and two statistically significant negative changes. BOD1L1 (FAM44A) is closely related to Bod1 (FAM44B) and Bod1L2 (FAM44C), but contains a short proline rich n-terminal extension, and a long, c-terminal extension (Figure 1H). Of note, one BOD1L1 peptide was doubly phosphorylated on T2956 and S2958 and displayed a significant increase (1.3-fold; *p* = 0.06) in phosphorylation in adapted cells relative to acutely treated cells. Another peptide was also doubly phosphorylated and contains S2973 and 2986 and displayed a significant increase (2.3-fold; *p <* 0.01) in phosphorylation in adapted cells relative to acutely treated cells. Another peptide was singly phosphorylated at S2986 and displayed a significant increase (1.2-fold; *p* = 0.07) in phosphorylation in adapted cells relative to acutely treated cells. Importantly, the total protein level of BOD1L1 did not change in adapted cells relative to acutely treated cells (1.06-fold; *p* = 0.26; 15 peptides) (Figure 1F, supplemental table 1) indicating that BOD1L1 undergoes net changes in phosphorylation status as cell adapt to UMK57. The peptide containing S2954, T2956 and S2958 was previously identified in a phospho-proteomic screen^14^, and the conserved cluster of lysine and arginine residues proximal to the phosphoacceptors sites suggest phosphorylation by Aurora kinases (figure 1I). Indeed, previous proteomic analysis demonstrated that phosphorylation of this peptide is sensitive to an Aurora kinase A inhibitor in a dose-dependent manner, and insensitive to an inhibitor of Aurora kinase B (Figure S1H)^14^. Moreover, BOD1L1 was previously detected in an Aurora kinase A proximity-ligation mapping experiment^15^. These data indicate that BOD1L1 is selectively phosphorylated by Aurora kinase A as cells adapt to UMK57. Interestingly, BOD1L1 is distinguished from other members of this gene family because neither Bod1 nor Bod1L2 contain obvious Aurora kinase consensus sites and the phosphoacceptors sites identified here and in previous proteomic screens are in the extended c-terminal domain that is unique to BOD1L1, in agreement with the fact that BOD1L1 is functionally distinct from BOD1 during DNA repair^16^.

### BOD1L1 and Aurora kinase A are required for adaptation to UMK57

To determine if BOD1L1 is functionally required for adaptive resistance to UMK57 in CIN cancer cells, we used two independent shRNA sequences to knockdown BOD1L1 expression and selected cells for shRNA expression using puromycin treatment. We confirmed a durable knockdown of BOD1L1 over several days in cells expressing BOD1L1 shRNA compared to cells expressing control shRNA using BodlL1-specific antibodies on immunoblot and through indirect immunofluorescence. (Figure S2A, S2B).

Next, we tested if BOD1L1 is required for cells to adapt to UMK57. We transfected cells with BOD1L1-specific shRNA or control shRNA expressing plasmids, selected for puromycin resistance, added UMK57 for short or prolonged periods and then measured the frequency of lagging chromosomes during mitosis. As expected in cells transfected with control shRNA, acute treatment with UMK57 reduces the frequency of lagging chromosomes in anaphase and high lagging chromosome rates return upon prolonged treatment with UMK57. Cells transfected with either of two distinct BOD1L1 shRNA’s display elevated rates of lagging chromosomes in anaphase compared to cells transfected with control shRNA. Despite the identified role for BOD1L1 in DNA repair, the increased rate of lagging chromosomes that we observe in BOD1L1 knockdown cells is not likely due to elevated levels of DNA damage, as it was previously demonstrated that there is no increase in basal levels of DNA repair foci in knockdown cells compared to control cells in the absence of prior cell irradiation^17^. BOD1L1-deficient cells respond to acute treatment with UMK57 and show significantly fewer anaphase cells with lagging chromosomes. The effect of UMK57 on BOD1L1-deficient cells is durable because cells retain significantly reduced lagging chromosome rates compared to DMSO-treated cells throughout prolonged treatment (Figure 2A, 2B). Therefore, BOD1L1 expression is required for adaptation to prolonged UMK57 treatment.

**Figure 2:**
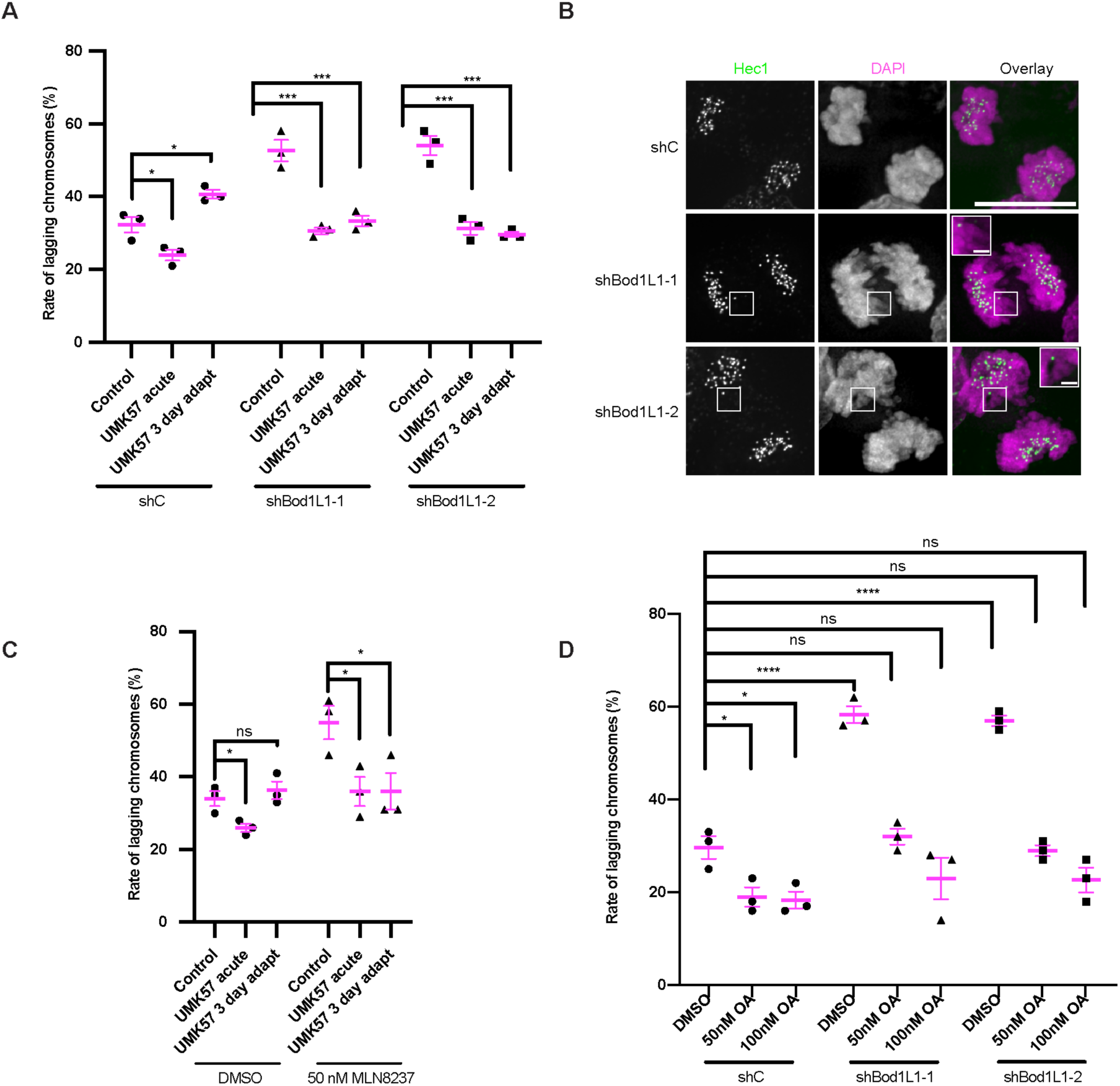
BOD1L1 and Aurora A are required for adaptation to UMK57. A) SW620 cells were transfected with control shRNA or shRNA against BOD1L1 and selected with puromycin. Following selection, the cells were fixed and stained for Hec1 and DAPI. Control cells were treated with DMSO and cells adapted to UMK57 were treated for 72 hours. Cells treated acutely with UMK57 were treated for 1 hour. The percentage of lagging chromosomes in anaphase for cells expressing the indicated shRNA is shown. 100 cells per shRNA and condition were scored for the presence of lagging chromosomes for each of 3 independent biological repeats. Error bars indicate the mean +/- SEM. * denotes *p* < 0.05, *** denotes *p* < 0.001. Statistical significance was calculated between the indicated conditions using Dunnett’s multiple comparison test. B) SW620 cells were transfected with control shRNA or shRNA against BOD1L1 and selected with puromycin. Following selection, the cells were fixed and stained for Hec1 and DAPI. Control cells were treated with DMSO, and cells adapted to UMK57 were treated for 72 hours. Cells treated acutely with UMK57 were treated for 1 hour. The cells were then fixed and stained for Hec1 and DAPI. Representative images from 3 independent experiments are shown. The images were adjusted evenly for brightness and contrast for presentation. The scale bars of main images are 10 μm, and insets are 1 μm. C) SW620 cells were plated. 24 hours following plating, DMSO or 50 nM MLN8237 was added, with or without 100 nM UMK57. After 72 hours, the cells were fixed and stained for Hec1 and DAPI. The percentage of lagging chromosomes in anaphase for cells expressing the indicated shRNA is shown. 100 cells per shRNA and condition were scored for the presence of lagging chromosomes for each of 3 independent biological repeats. Error bars indicate the mean +/- SEM. * denotes *p* < 0.05, *** denotes *p* < 0.001. Statistical significance was calculated between the indicated conditions using Dunnett’s multiple comparison test. D) SW620 cells were transfected with control shRNA or shRNA against BOD1L1 and selected with puromycin. Following selection, the cells were treated with DMSO or Okadaic acid as indicated for 90 minutes prior to fixation. The cells were then stained for Hec1 and DAPI. The percentage of lagging chromosomes in anaphase for cells expressing the indicated shRNA is shown. 100 cells per shRNA and condition were scored for the presence of lagging chromosomes for each of 3 independent biological repeats. Error bars indicate the mean +/- SEM. * denotes *p* < 0.05, *** denotes *p* < 0.001. Statistical significance was calculated between the indicated conditions using Dunnett’s multiple comparison test.

Since BOD1L1 is phosphorylated by Aurora kinase A in mitosis and there was a significant increase in BOD1L1 phosphorylation during adaptation to prolonged UMK57 treatment, we also tested if Aurora kinase A activity is required for adaptation to UMK57. We treated cells concurrently with a sublethal 50 nM dose of Aurora kinase A inhibitor and UMK57 over 3 days and assessed the rates of lagging chromosomes. SW620 cells continued to proliferate in the presence of this dose of Aurora kinase A inhibitor (Figure S2C), and cells treated with Aurora kinase A inhibitor alone displayed elevated levels of lagging chromosomes compared to cells treated with DMSO. Cells treated with Aurora kinase A inhibitor respond to acute treatment with UMK57 and display significantly fewer lagging chromosomes in anaphase relative to cells treated with the Aurora kinase A inhibitor alone. The fraction of anaphase cells with lagging chromosomes remained significantly reduced in cells with prolonged UMK57 treatment when Aurora kinase A was inhibited demonstrating that Aurora kinase A is also required for adaptation to prolonged UMK57 treatment (Figure 2C).

These and previous data^11^ indicate that adaptation to prolonged UMK57 treatment likely occurs through changes in kinetochore protein phosphorylation and that BOD1L1 may play a role in regulating protein phosphorylation. Therefore, we tested if treatment of cells with a modest dose of Okadaic acid^18^, which inhibits PP2A and to a lesser extent PP1 phosphatases, would alter the rate of lagging chromosomes in anaphase. Cells transfected with control shRNA displayed a reduced frequency of lagging chromosomes in anaphase when treated with Okadaic acid, suggesting that elevated protein phosphorylation can increase chromosome segregation fidelity (Figure 2D). As expected, BOD1L1-deficient cells display elevated numbers of lagging chromosomes in anaphase. Remarkably, cells transfected with shRNA against BOD1L1 and treated with Okadaic acid had rates of lagging chromosomes in anaphase that were not different from control shRNA/DMSO treated cells. Thus, Okadaic acid treatment effectively replaces BOD1L1 function in mitosis as measured by chromosome segregation fidelity indicating that BOD1L1 likely acts as a negative regulator of protein phosphatase activity and that the elevated rate of lagging chromosomes in BOD1L1-deficient cells (Figure 2A) is due to reduced levels of protein phosphorylation.

Since Aurora kinase A is required for adaptation to UMK57, we used cell biological approaches to test if Aurora kinase A activity is altered in cells treated either acutely with or adapted to UMK57. We measured the intensity of Aurora kinase A and pT288 Aurora kinase A, a marker of Aurora kinase A activity in mitotic cells from an asynchronous culture. Whereas the ratio of Aurora kinase A to pT288 Aurora kinase A remained unchanged in cells treated acutely with or adapted to UMK57 compared to control cells (i.e., no change in phospho-occupancy), the overall levels of both Aurora kinase A and pT288 Aurora kinase A on mitotic spindles strongly increased in cells treated with UMK57 (Figure 3A, 3B). Since TPX2 directs Aurora kinase A to the mitotic spindle^19^, we quantified the levels of TPX2 and alpha tubulin in the mitotic spindle in cells treated with UMK57. Similar to Aurora kinase A, this experiment revealed elevated levels of TPX2 on the mitotic spindle in cells treated with UMK57 despite a decrease in overall levels of spindle microtubule content in treated cells (Figure 3C, 3E). TACC3, another spindle-associated protein, displayed spindle intensity that roughly mirrored the levels of spindle microtubule content (Figure 3D, 3E) indicating that the quantity of Aurora kinase A and TPX2 on spindles selectively increased upon UMK57 treatment.

**Figure 3:**
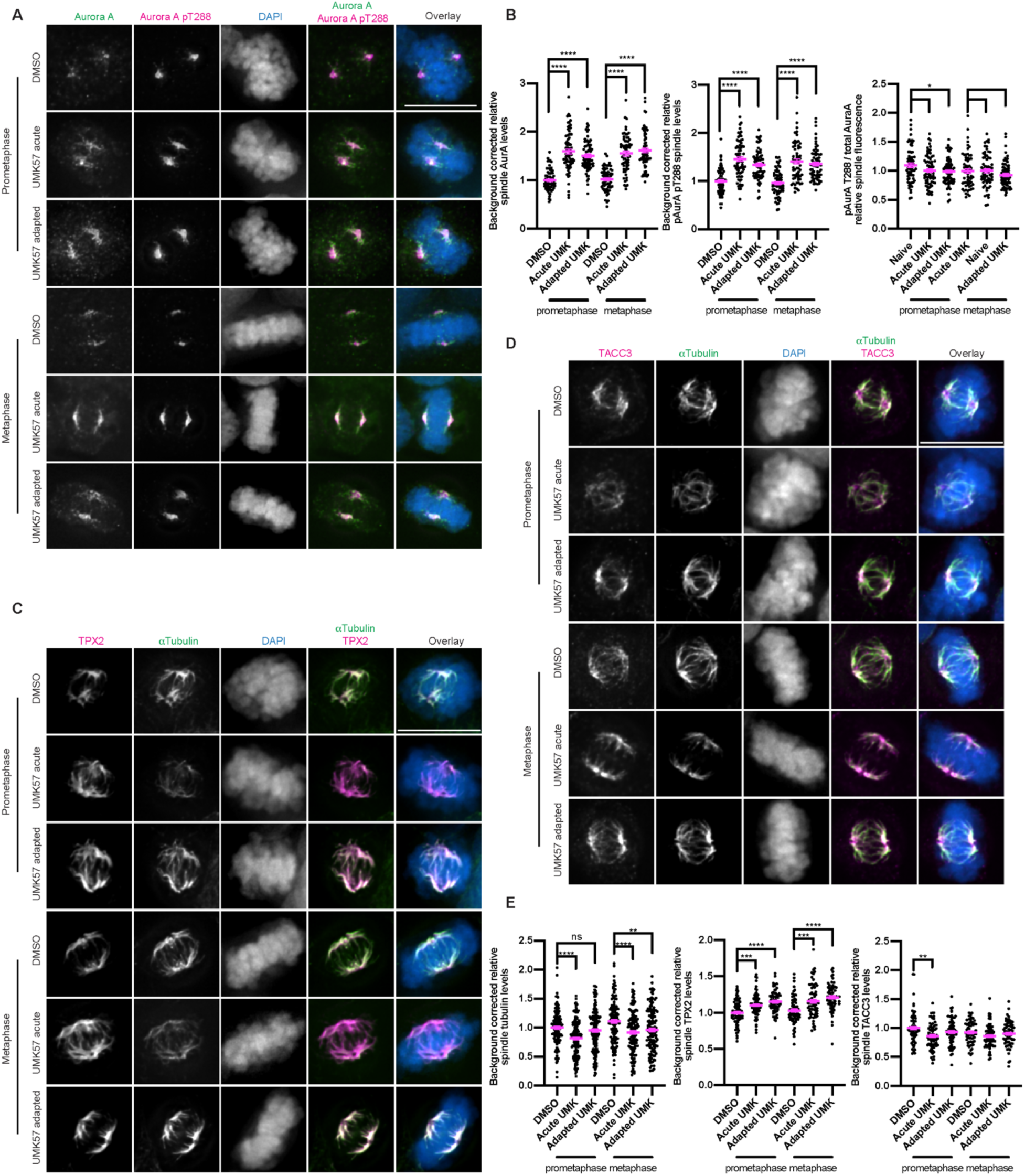
Microtubule destabilization mediated by hyperactive MCAK enhances Aurora A/TPX2 localization to the mitotic spindle. A) Immunofluorescence images from an asynchronous population of SW620 cells treated either with DMSO control, acute (1 hour) UMK57 or UMK57 for 3 days. The cells were then fixed and stained for total Aurora A and Aurora A pT288. The Aurora A and Aurora A pT288 were adjusted evenly for brightness and contrast for presentation. The DAPI channel of each condition was adjusted independently. Representative images from 2 independent biological experiments are shown. The scale bars are 10 μm. B) Quantification of the relative spindle protein intensities from the conditions from panel (A). The levels of the prometaphase DMSO conditions was set to 1 and the other conditions shown as fold-changes. The fluorescence levels for at least 30 cells for each of 2 independent biological repeats were measured. Error bars indicate the mean +/- SEM. Statistical significance was calculated between the indicated conditions using Dunnett’s multiple comparison test. C) Cells were prepared as for panel (A) but stained for TPX2 and alpha tubulin. D) Cells were prepared as for panel (A) but stained for TACC3 and tubulin. E) Quantification of the relative spindle protein intensities from panels (C) and (D). The tubulin intensities from panels (C) and (D) were combined into a single plot.

### BOD1L1 regulates mitotic progression and mitotic fidelity

To date, the only role identified for BOD1L1 is in DNA repair^16,17,20^. BOD1L1 shares extensive sequence similarity to Bod1 which has been shown to be involved in regulation of mitotic activities including kinetochore protein phosphorylation by recruiting the PP2A-B56 phosphatase, regulating MCAK localization, and also to be important for chromosome alignment^21–23^. Given the similarities between Bod1 and BOD1L1, and that BOD1L1-deficient cells display increased rates of lagging chromosomes in anaphase and that BOD1L1 function in mitotic fidelity can be rescued using protein phosphatase inhibitors, we focused our efforts on understanding the contribution of BOD1L1 to mitotic processes and chromosome segregation fidelity.

Using immunofluorescence microscopy we detected BOD1L1 at spindle poles, on microtubules and at kinetochores of SW620 cells (Figure 4A, 4B). Line scans showed significant overlap between BOD1L1 and Hec1 at kinetochores in prometaphase through anaphase. Notably, BOD1L1 does not localize to all kinetochores, but rather to a subset, suggesting the possibility of regulation of BOD1L1 localization. Since BOD1L1 localizes to the mitotic spindle, we also examined if BOD1L1 co-localizes with Aurora kinase A. We found a large degree of co-localization between Aurora kinase A and BOD1L1 at the spindle poles and along pole-proximal microtubules (Figure 4C). BOD1L1 co-localized with Hec1 on kinetochores in cells treated with either a low or high dose of nocodazole demonstrating that BOD1L1 does not require microtubules for kinetochore localization (Figure S2D). BOD1L1 localization was also unchanged by treatment of cells with Aurora kinase A or Aurora kinase B inhibitors demonstrating that BOD1L1 kinetochore localization is independent of these kinase activities despite the problems in spindle structure and chromosome alignment caused by these inhibitors (Figure S2E).

**Figure 4:**
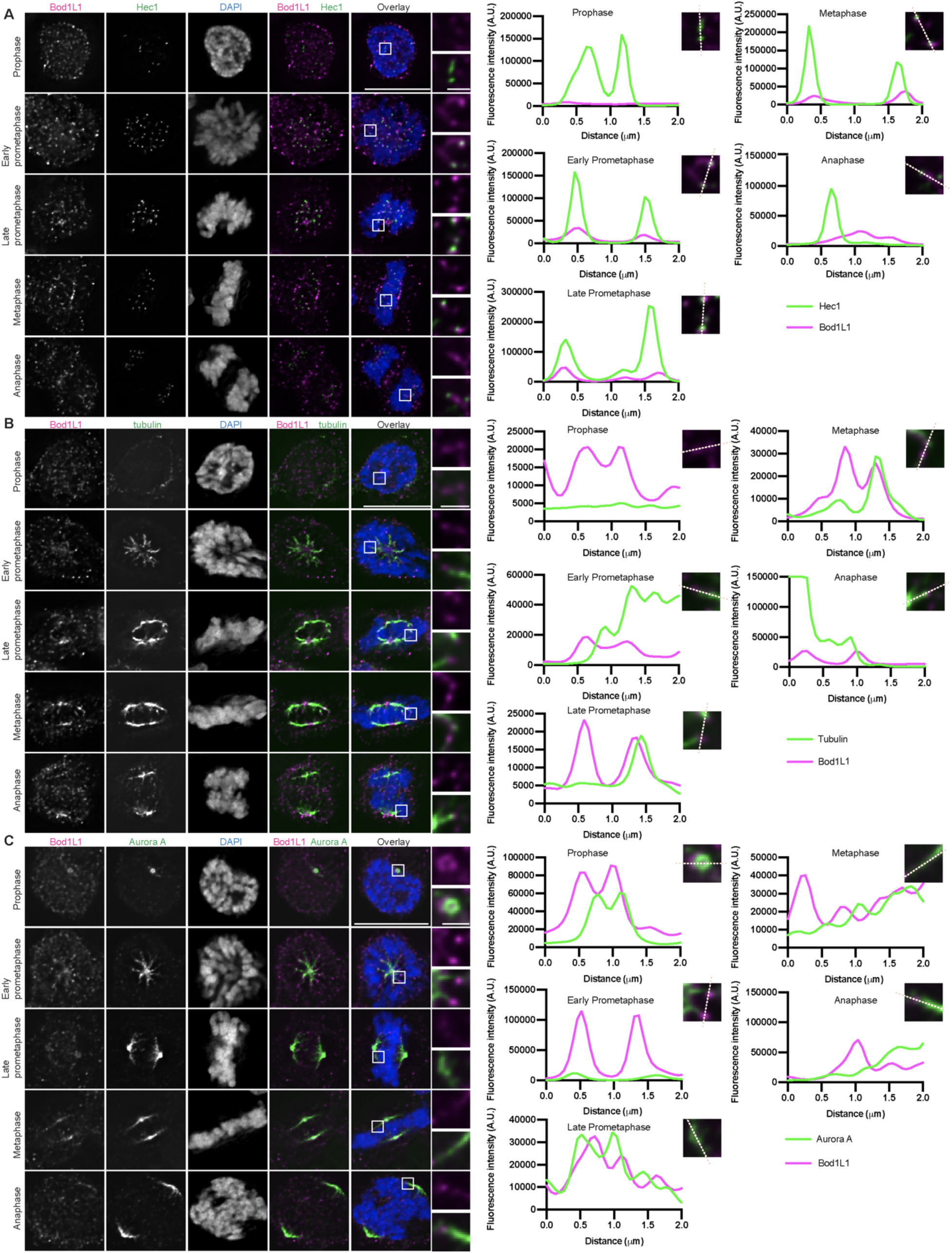
BOD1L1 localizes to the mitotic spindle and kinetochores during mitosis. A) Immunofluorescence images from an asynchronous population of SW620 cells in various stages of mitosis stained for BOD1L1, Hec1 and DAPI. The BOD1L1 and Hec1 channels were adjusted evenly for brightness and contrast for presentation. The DAPI channel of each condition was adjusted independently. Representative images from 2 independent biological experiments are shown. The scale bars of main images are 10 μm, and insets are 1 μm. B) Immunofluorescence images from an asynchronous population of SW620 cells in various stages of mitosis stained for BOD1L1, alpha tubulin and DAPI. The BOD1L1 and alpha tubulin channels were adjusted evenly for brightness and contrast for presentation. The DAPI channel of each condition was adjusted independently. Representative images from 2 independent biological experiments are shown. The scale bars of main images are 10 μm, and insets are 1 μm. C) Immunofluorescence images from an asynchronous population of SW620 cells in various stages of mitosis stained for BOD1L1, Aurora A and DAPI. The BOD1L1 and Hec1 channels were adjusted evenly for brightness and contrast for presentation. The DAPI channel of each condition was adjusted independently. Representative images from 2 independent biological experiments are shown. The scale bars of main images are 10 μm, and insets are 1 μm.

BOD1L1-deficient cells appear to have aberrant spindle structures and problems with chromosome alignment. We quantified these phenotypes by measuring the proportion of cells in each phase of mitosis as a percentage of all mitotic cells. BOD1L1-deficient early prometaphase cells often had small spindles with kinetochores located close to the poles, whereas late prometaphase cells had difficulties in chromosome alignment (Figure 5A, 5B), with many chromosomes located proximal to the spindle poles. We also used time-lapse microscopy on mCherry-H2B expressing SW620 cells to measure the time spent in mitosis with and without BOD1L1. Control cells needed on average 57 minutes to transit mitosis from the onset of chromosome condensation to anaphase. In contrast, BOD1L1-deficient cells spent on average 208.3 minutes (shRNA #1) and 120.6 minutes (shRNA #2) in mitosis. These changes in the average time taken to complete mitosis overestimates the overall impact of mitotic delay in BOD1L1-deficient cells because it is skewed by cells that either underwent slippage or death following extended times in mitosis. Amongst cells that completed mitosis with anaphase, these cells took an average of 137.5 (shRNA #1) and 100.0 minutes (shRNA #2) to transit mitosis (Figure 5C, S3A). Thus, BOD1L1 is functionally required for efficient mitotic progression.

**Figure 5:**
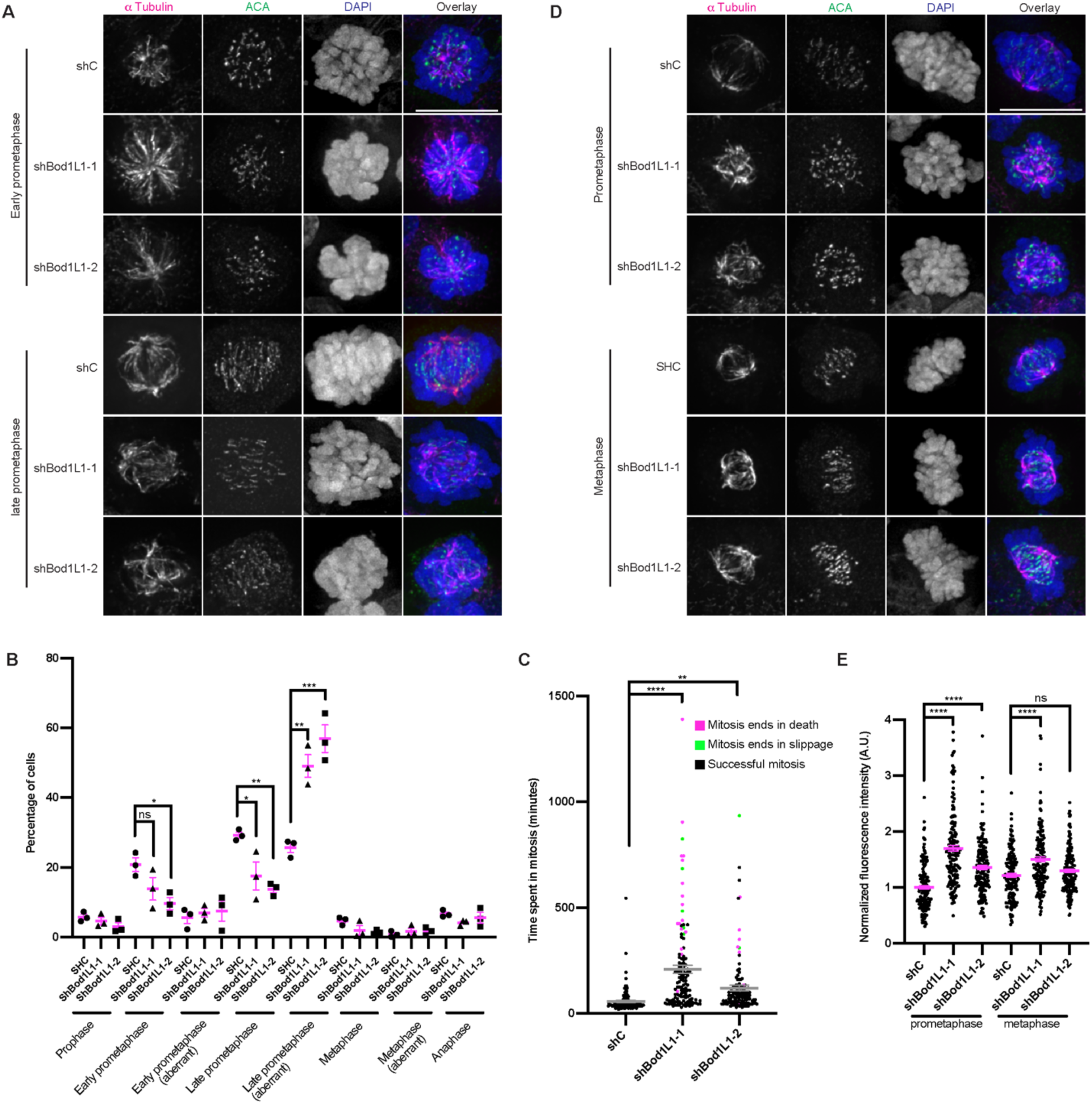
BOD1L1 is required for correct k-MT attachment stability and faithful chromosome segregation. A) Immunofluorescence images of SW620 cells were transfected with control shRNA or shRNA against BOD1L1 and selected with puromycin. The cells were then fixed and stained for alpha tubulin and ACA. Representative images of 2 independent biological experiments are shown. The images were adjusted evenly for brightness and contrast for presentation. The scale bar is 10μm. B) Quantification of the cells shown in panel (A). Statistical significance was calculated between the indicated conditions using Dunnett’s multiple comparison test. C) SW620 cells were co-transfected with control shRNA or shRNA against BOD1L1 together with mCherry-H2B and selected with puromycin. Following selection, time lapse imaging was started at 5-minute intervals. The fate of at least 40 cells per experiment for each of 3 independent biological repeats was recorded. Statistical significance was calculated between the indicated conditions using Dunnett’s multiple comparison test. D) Immunofluorescence images of SW620 cells were transfected with control shRNA or shRNA against BOD1L1 and selected with puromycin. The cells were then placed at 4°C for 15 minutes before they were fixed and stained for alpha tubulin and ACA. Representative images of 3 independent biological experiments are shown. The images were adjusted evenly for brightness and contrast for presentation. The scale bar is 10μm. E) Quantification of the relative intensity of alpha Tubulin from the conditions shown in panel (D). The levels of prometaphase shC expressing cells were set to 1 and the other conditions shown as a fold-change. At least 50 cells per condition from each of 3 independent biological experiments were quantified. Error bars indicate the mean +/- SEM. **** denotes *p* < 0.0001. Statistical significance was calculated between the indicated conditions using Dunnett’s multiple comparison test.

Next, we tested if BOD1L1 contributes to k-MT attachment stability. Following transfection with control or BOD1L1-specific shRNA plasmids and puromycin selection, we cold treated cells to eliminate non-k-MT from spindles and fixed and stained cells for tubulin and ACA. The spindle microtubule intensity in BOD1L1-deficient cells was increased in both prometaphase and metaphase relative to cells transfected with control shRNA (Figure 5D, E).

### BOD1L1 interacts with the PP2A-B56 phosphatase and regulates kinetochore phosphorylation

To gain further insight into the role(s) of BOD1L1 in mitosis, we immunoprecipitated BOD1L1 from mitotic cells to determine the identity of interacting proteins. We cross-linked antibody to protein G beads and used them to immunoprecipitate BOD1L1 interacting proteins from SW620 cells synchronized in mitosis through sequential treatment with thymidine and nocodazole. The eluted proteins were then separated by SDS-PAGE gel, stained with Coomassie, cut into fractions and analyzed by mass spectrometry (Figure 6A). This analysis revealed a high affinity interaction with the MTORC2 signaling complex which regulates cellular pro-survival and is typified by the presence of the Rictor protein subunit^24^ as judged by the very high number of detected peptides. Also, there were numerous mRNA translation initiation factor proteins, and abundant peptides from the CCT chaperone complex. In addition, we detected several proteins that participate in DNA repair, including SETD1A, RAD50, XRCC5, XRCC6, RIF1 and MRE11, consistent with previous findings, and which serve as positive controls for the specificity of our assay^17,20^. We also detected various core subunits of the Anaphase Promoting Complex/Cyclosome, although we did not detect the co-activator proteins Cdh1 or Cdc20. Thus, BOD1L1 may be an APC/C substrate or a regulator of APC/C activity. We also detected numerous proteins with documented mitotic functions including Sgo2, CENPF, MCAK, Numa, Aurora kinase A, and several components of the PP2A-B56 phosphatase complex including both PPP2R1A and PPP2R1B regulatory subunits, the PPP2CA catalytic subunit, and the PPP2R5E epsilon adaptor subunit (Figure 6B, Supplemental table 3). Therefore, like Bod1, BOD1L1 interacts with PP2A and likely regulates PP2A activity or is itself regulated by PP2A. These interacting proteins were independently validated by blotting for various proteins. We detected the PP2A catalytic, regulatory and the delta, gamma, and epsilon adaptor subunits, but not the beta subunit. We also detected CENPF, MCAK, Bub3, SMC1 and APC3 proteins in BOD1L1 immunoprecipitates (Figure 6C). These interactions were generally enriched when BOD1L1 was immunoprecipitated from cells synchronized in mitosis, compared to asynchronous cells.

**Figure 6:**
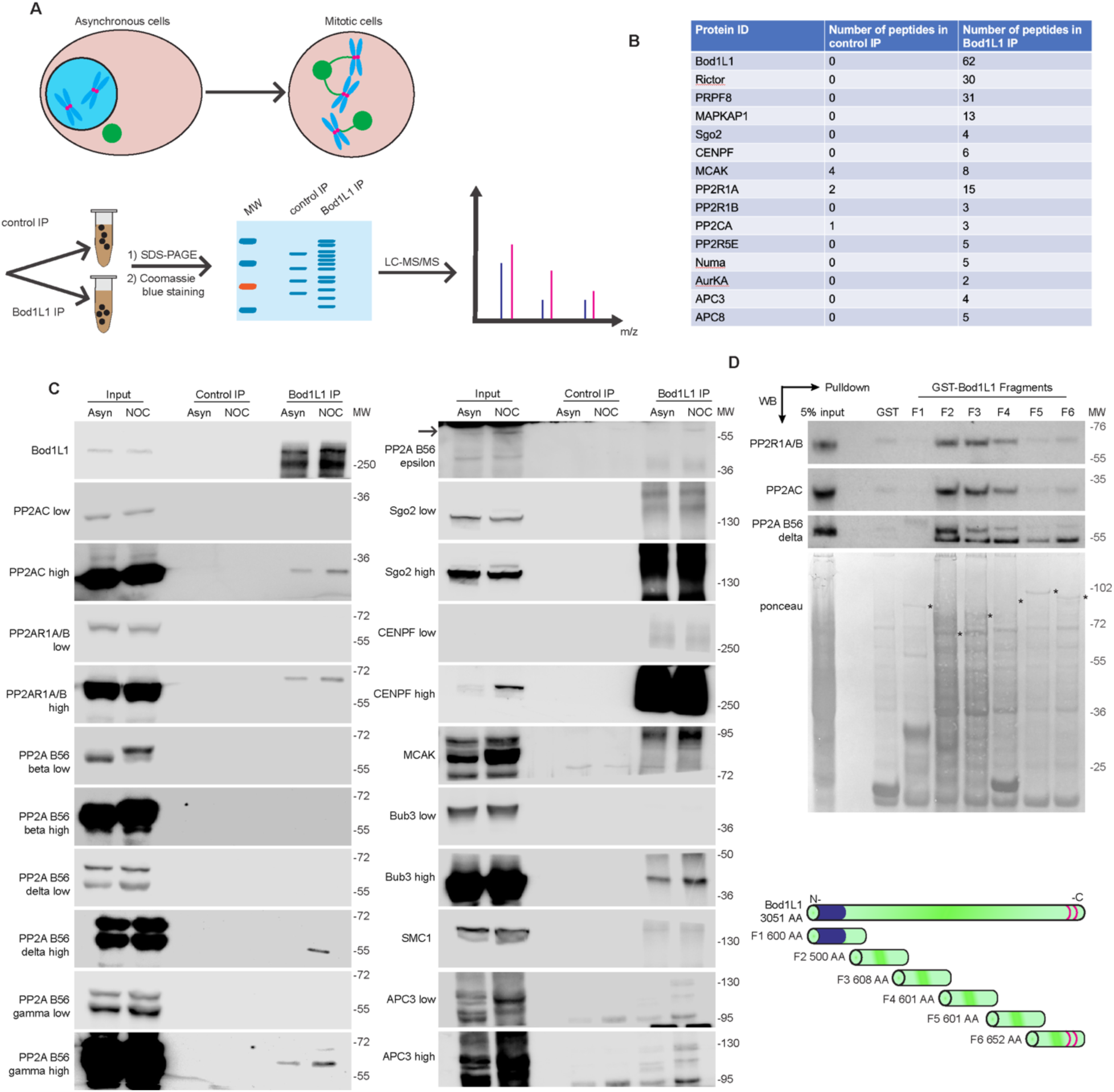
BOD1L1 interacts with the PP2A-B56 phosphatase complex. A) Cartoon illustration depicting the experimental scheme used to discover protein interactors of BOD1L1. SW620 cells were first synchronized by thymidine-nocodazole. BOD1L1 was then immunoprecipitated and purified along with interacting proteins. The samples were then separated by SDS-PAGE. The entire lanes were excised and cut into 5 pieces and analyzed by mass spectrometry. The experiment was performed once. B) Table of selected interacting proteins detected in the control immunoprecipitation and BOD1L1 immunoprecipitations showing the number of detected peptides. C) SW620 cells were prepared as in panel (A), except that they were analyzed by Western blot. BOD1L1 was immunoprecipitated and the purified proteins were then separated by SDS-PAGE, transferred to nitrocellulose membrane and blotted as indicated. The panels were adjusted for brightness and contrast for presentation. An experiment representative of two independent biological repeats is shown. D) SW620 mitotic nuclear cell extracts were incubated with GST or GST-BOD1L fragments. Complexes were then isolated by glutathione-sepharose and analyzed by Western blot. The blot shown is representative of 2 independent experiments.

There are notable similarities and differences in the interactomes of Bod1 and BOD1L1^23^. Like Bod1, BOD1L1 interacts with PP2A-B56, SMC1A, SMC3, SETD1A, Aurora kinase A and Bub3. In contrast, Bod1 did not show interaction with the MTORC2 complex, but does interact with Hec1, Knl1 and components of the fibrous corona including Zwint, ZW10 and dynein. Hec1 is among the strongest interacting proteins with Bod1 but was not detected as an interactor with BOD1L1 in our analysis despite the co-localization of BOD1L1 with Hec1 at kinetochores at the resolution of immunofluorescence microscopy. To test directly if BOD1L1 kinetochore localization requires Hec1, we utilized HeLa cells with doxycycline inducible CRISPR/Cas9 Hec1 genetic knockout^25^, and compared the localization of BOD1L1 in the presence and absence of Hec1. We did not detect a significant change in BOD1L1 localization indicating that Hec1 is not required for BOD1L1 kinetochore localization (Figure S3B).

To determine which domain of BOD1L1 is responsible for interaction with the PP2A complex, we performed a pull-down experiment using fragments of BOD1L1 fused to GST as bait from lysates of mitotic SW620 cells. Surprisingly, despite the high degree of similarity between the n-terminus of BOD1L1 and Bod1 and the fact that both Bod1 and BOD1L1 interact with PP2A subunits, the region of BOD1L1 that bound to the PP2A complex most strongly was the uncharacterized central domain from amino acid 500 to 2001. Additionally, the c-terminal F6 fragment also interacted with the PP2A B56 delta subunit (Figure 6D). In contrast, it is the n-terminal F1-F2 Bod1-similar and C-terminal F6 regions that bind Rif1 in the context of the DNA damage response^17^.

These data indicate that BOD1L1 might regulate the PP2A-B56 complex during mitosis to regulate mitotic fidelity. Therefore, we tested whether BOD1L1 regulates kinetochore protein phosphorylation levels in a similar manner to Bod1. Using immunofluorescence, we quantified the staining intensity for numerous phosphorylated proteins at kinetochores following depletion of BOD1L1. We stained cells for Aurora kinase A pT288 (Aurora kinase A), Hec1 pS55 (Aurora kinase A), Hec1 pT31 (Cdk1-Cyclin B1), Hec1 pS44 (Aurora kinase B), Dsn1 pS100 (Aurora kinase B), Knl1 pMELT (Mps1), Aurora kinase B pT232 (Aurora kinase B) and MCAK pS95 (Aurora kinase B). We observed that the intensity of staining for Hec1 pS55, Hec1 pT31, Hec1 pS44, pDsn1 pS100, and Knl1 pMELT decreased in both prometaphase and metaphase and Aurora kinase B pT232 levels decreased in prometaphase in BOD1L1-deficient cells. In contrast, MCAK pS95 staining intensity increased in metaphase cells (Figure 7). Notably, we did not detect a pattern of changes for Aurora kinase A pT288, indicating that total Aurora kinase A activity levels are essentially unchanged in BOD1L1-deficient mitotic cells. To verify that the observed changes in kinetochore protein phosphorylation were directly caused by loss of BOD1L1, we transfected SW620 BOD1L1-knockdown cells with a plasmid to express GFP-BOD1L1 and quantified the level of Hec1-pT31 and KNL1-pMELT during mitosis. There is a detectable increase in BOD1L1 expression in cells expressing GFP-BOD1L1 although the molecular weight change of the fusion protein is essentially unresolvable from endogenous because of the very large molecular weight of BOD1L1 (>330k). The expression of GFP-Bod1L1 restored Hec1 an KNL1 phosphorylation levels on these phosphoacceptors sites in the knockdown cells and slightly increased the relative phosphorylation levels compared to control cells (Figure S4A,B,C,F,G). Furthermore, to confirm BOD1L1 mitotic function in non-transformed cells, we knocked down BOD1L1 with two independent shRNA sequences in the non-tranformed and diploid cell line RPE1 and quantified the level of Hec1-pT31 and KNL1-pMELT during mitosis. Transfection with either shRNA sequence induced a significant reduction in the relative phosphorylation levels of Hec1 and KNL1 in RPE1 cells compared to control cells consistent with observations in the transformed SW620 cells (Figure S4D,E,H,I). These results combined with previous data showing that phosphatase inhibitors can rescue lagging chromosomes in BOD1L1-deficient mitotic cells demonstrate that BOD1L1 functions as a negative regulator of protein phosphatase activity to support efficient mitotic progression. Consistently, the loss of BOD1L1 affects phospho-substrates at kinetochores that are phosphorylated by multiple distinct protein kinases.

**Figure 7:**
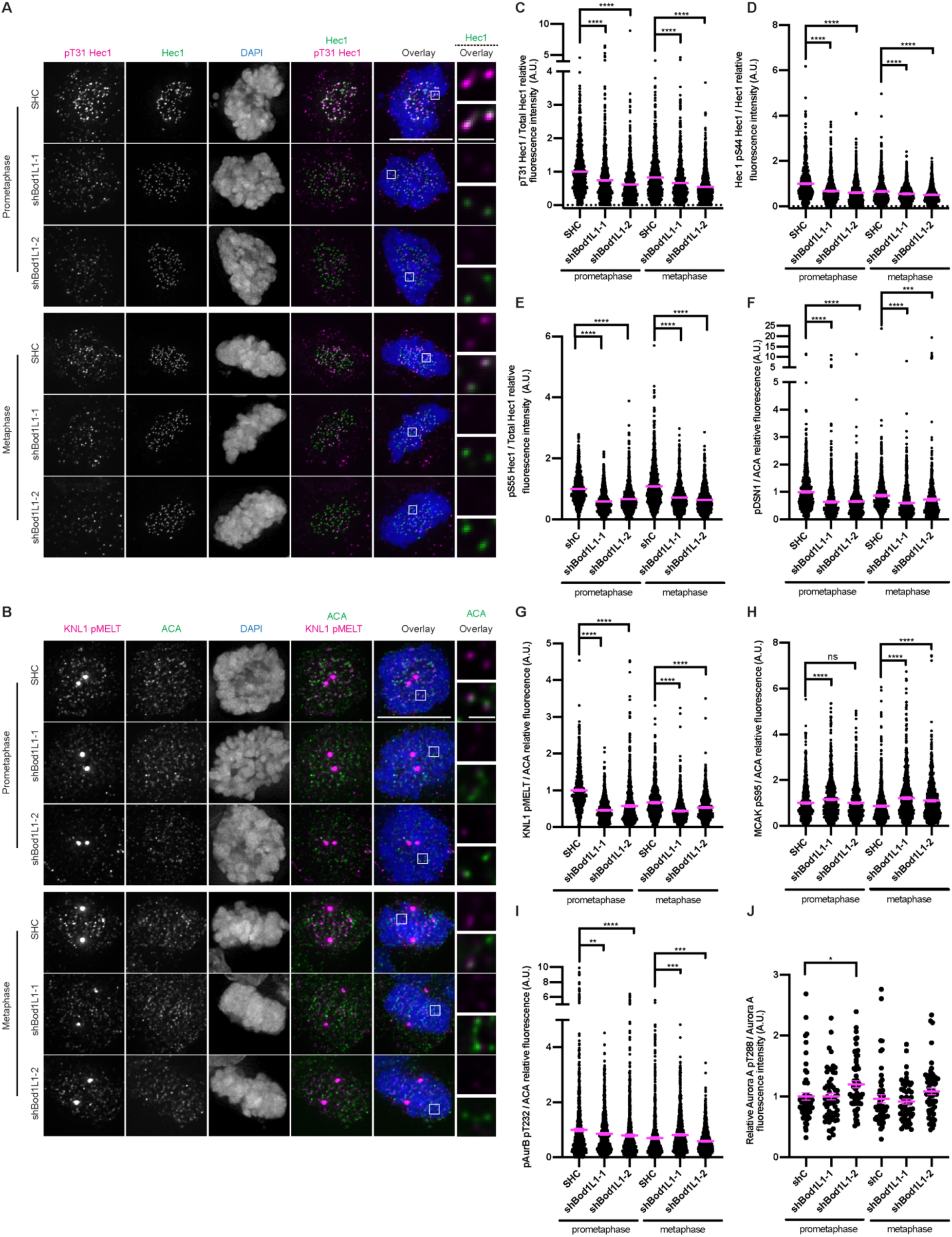
BOD1L1 expression is required for normal kinetochore phosphorylation. A) Immunofluorescence images from an asynchronous population of SW620 cells transfected with control shRNA or shRNA against BOD1L1 in prometaphase or metaphase and stained for Hec1, Hec1 pT31 and DAPI. The BOD1L1 and Hec1 channels were adjusted evenly for brightness and contrast for presentation. The DAPI channel of each condition was adjusted independently. Representative images from 2 independent biological experiments are shown. The scale bars of main images are 10 μm, and insets are 1 μm. B) As for panel (A), except for KNL1 pMELT/ACA instead of Hec1 pT31/total Hec1 C) Quantification of the relative kinetochore pT31 Hec1/Hec1 intensities from the conditions from panel (A). The condition with the lowest level of pT31 Hec1/Hec1 was set to 1 and the other conditions shown as fold-changes. 25 kinetochores were quantified from each of 20 cells for each of 2 independent biological repeats. Error bars indicate the mean +/- SEM. Statistical significance was calculated between the indicated conditions using Dunnett’s multiple comparison test. D) As for panel (C), except for Hec1 pS44/total Hec1instead of Hec1 pT31/total Hec1 E) As for panel (C), except for Hec1 pS55/total Hec1 instead of Hec1 pT31/total Hec1 F) As for panel (C), except for DSN1 pS109/ACA instead of Hec1 pT31/total Hec1 G) As for panel (C), except for KNL1 pMELT/ACA instead of Hec1 pT31/total Hec1 H) As for panel (C), except for MCAK pS95/ACA instead of Hec1 pT31/total Hec1 I) As for panel (C), except for Aurora B pT232/ACA instead of Hec1 pT31/total Hec1 J) Quantification of total cellular Aurora A pT288/Total Aurora A. The condition with the lowest level of pT31 Hec1/Hec1 was set to 1 and the other conditions shown as fold-changes. The level of protein was measured from at least 25 cells for each of 2 independent biological repeats. Error bars indicate the mean +/- SEM. Statistical significance was calculated between the indicated conditions using Dunnett’s multiple comparison test.

### *BOD1L1* gene is mutated in human cancers

Since aneuploidy is associated with cancer development and BOD1L1-deficiency increases mitotic error rates, we examined whether BOD1L1 is mutated in human cancers. We queried The Cancer Genome Atlas (TCGA) for cancers that have somatic alterations in the wild-type sequence in *BOD1L1*. Among the 10967 cancers that have been sequenced, 3.9% of cancers show alterations in *BOD1L1* sequence, although the prevalence of *BOD1L1* alterations was variable across cancer types. The five cancers with the highest percentage of *BOD1L1* somatic alterations are uterine endometrial carcinoma (12.9%), lung adenocarcinoma (8.8%), skin cutaneous melanoma (7.9%), stomach adenocarcinoma (7.7%) and lung squamous cell carcinoma (6.6%). Because our data demonstrate that BOD1L1 is activated by Aurora kinase A and functionally regulates PP2A activity, we also examined the TCGA for cancers that have alterations in any of *BOD1L1*, *AURKA, TPX2, PPP2CA, PPP2CB, PPP2R1A, PPP2R1B, PPP2R5A, PPP2R5B, PPP2R5C, PPP2R5D, PPP2R5E*. Surprisingly, 59.6% of uterine carcinosarcoma and 43.9% of uterine endometrial cancers have alterations in one or more of these genes, suggesting that genes in this biochemical pathway might be particularly relevant in these cancer types (Figure S5A). We next tested associations of *BOD1L1* alteration with 5-year overall survival adjusting for potential confounders in these patients. *BOD1L1* alteration was significantly associated with improved 5-year survival in uterine cancer HR = 0.37 (95% CI:0.16 – 0.85). *BOD1L1* alteration was significantly associated with reduced 5-year survival in lung adenocarcinoma HR = 1.66 (95% CI: 1.03-2.66) and in melanoma HR = 1.79 (95% CI: 1.07-3.00). *BOD1L1* alteration did not demonstrate a significant association with patient survival in stomach adenocarcinoma or lung squamous cell carcinoma (Figure 8B).

**Figure 8:**
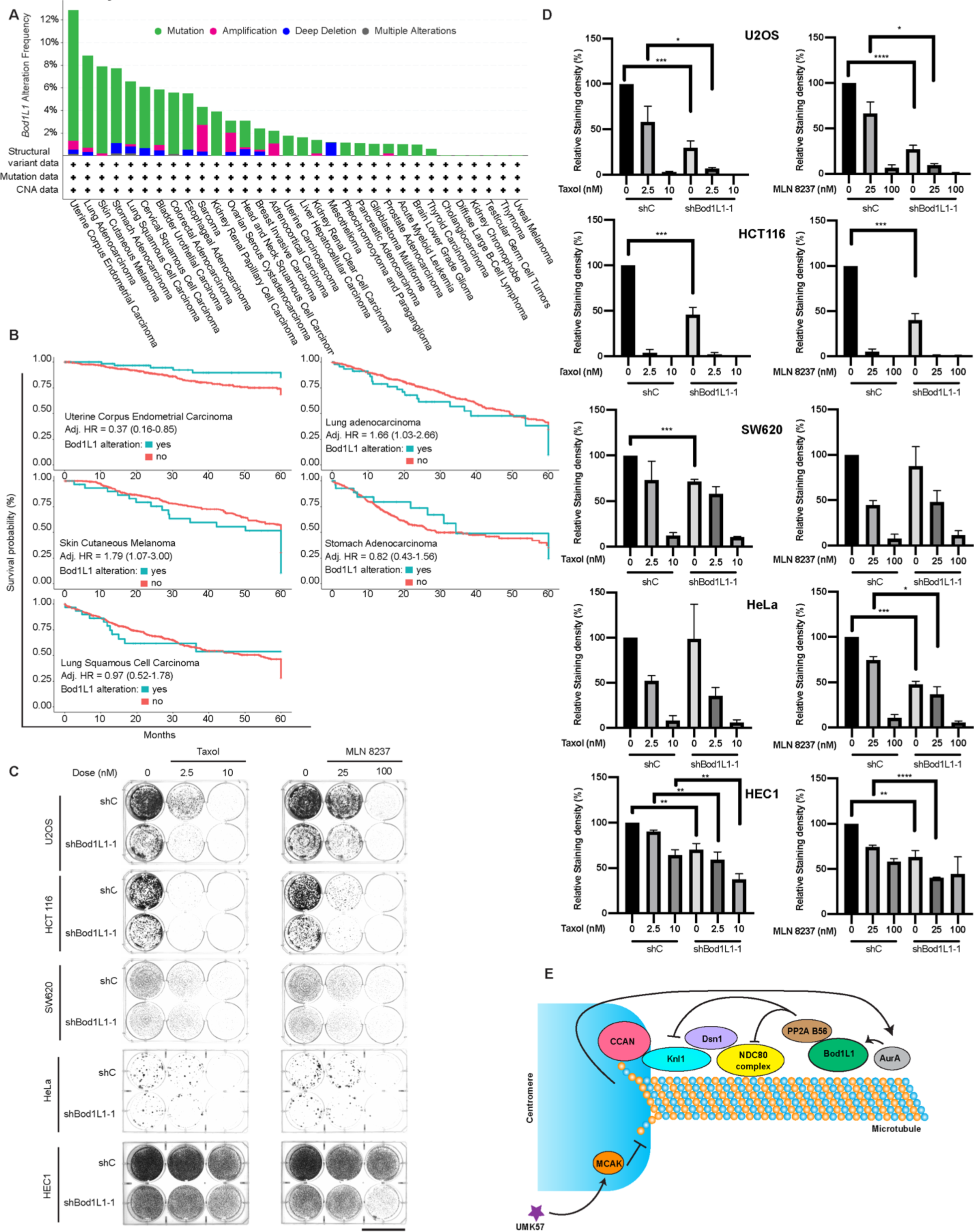
BOD1L1 is mutated in human cancers. A) The 10,967 sequenced cancers in the TCGA were queried for the presence of mutations in the *BOD1L1* gene and plotted by cancer subtype. B) Kaplan-Meier five-year overall survival by *BOD1L1* alteration status with Cox-Proportional Hazards Ratios and 95% confidence intervals from models adjusted for potential confounders (age, sex [except uterine], and tumor grade or stage). A) Uterine corpus endometrial carcinoma TCGA-UCEC, n= 529; b) lung adenocarcinoma TCGA-LUAD, n = 667; c) cutaneous melanoma TCGA-SKCM, n = 448; d) stomach adenocarcinoma TCGA-STAD, n = 440; e) lung squamous cell carcinoma TCGA-LUSC, n = 487. C) U2OS, HCT116, SW620 and HeLa cells were plated. They were then transfected with control shRNA or shRNA against BOD1L1 and selected with puromycin in the presence of compounds as indicated. After 1 week of selection, the cells were fixed and stained with crystal violet. Representative images from 3 independent experiments are shown. The scale bar is 4 cm. D) Quantification of cells from (C). The staining intensity of the cells was measured using Fiji and plotted. Error bars indicate the mean +/- SEM. Statistical significance was calculated between the indicated conditions using Student’s *t-* test. E) Cartoon illustration showing a possible mechanism for adaptation to UMK57. UMK57 agonizes MCAK, resulting in lower k-MT stability, which is sensed by Aurora A. Aurora A then phosphorylates BOD1L1, releasing PP2A to reduce phosphorylation of the NDC80 complex, resulting in higher k-MT stability.

Given that BOD1L1 appears to function in a pathway consisting of Aurora kinase A-BOD1L1-PP2A B56, we asked if individuals with mutations in *BOD1L1* also have co-occurring mutations in either *AURKA* or its cofactor *TPX2* or in *PP2A* complex components. Using data from the TCGA-PanCancer Atlas (n = 10,967), we observed that *BOD1L1* alterations were exclusive of alterations to other *AURKA-BOD1L1-PP2A* gene pathway members. This was true when testing *BOD1L1* alteration versus alteration to any other gene pathway member (Fisher’s exact P=1.3E-24), and *BOD1L1* alteration versus each of the eleven gene pathway members independently (all Fisher’s exact P<5.7E-08) (Figure S5B, S5C), suggesting that mutations in multiple components of this pathway are not tolerated in cancers.

To understand how BOD1L1 inactivation affects cell proliferation and response to anti-mitotic drugs including Aurora kinase A inhibitors or taxol, we compared the proliferation of cells derived from various tumor types [U2OS (osteosarcoma), HCT116 (colorectal carcinoma), SW620 (colorectal adenocarcinoma), HeLa (cervical adenocarcinoma) and HEC1 (uterine endometrial adenocarcinoma)] following transfection with either control shRNA or BOD1L1-specific shRNA. All of these cell lines express detectable BOD1L1, albeit at different levels, and they all require BOD1L1 for mitotic fidelity because there was a significant increase in lagging chromosome rates in all cell lines following shRNA-mediated depletion of BOD1L1. (Figure S5D & S5E). All cell types tested have reduced proliferation following transfection with the BOD1L1-specific shRNA compared to the control shRNA, although the magnitude of the reduction varied between cell types. This variance was not proportional to the level of BOD1L1 expression (e.g. HEC1 and SW620 cells have comparable expression levels, but different BOD1L1 sensitivities) (Figure 8C, 8D, S5D, S5G). The proliferation of these cells following treatment with taxol and Aurora kinase A inhibitor also varied. Proliferation of HCT116 cells was modestly reduced by depletion of BOD1L1, but profoundly suppressed by treatment with either dose of either drug with or without BOD1L1. In contrast, there was a dose-dependent reduction in proliferation of U2OS, HeLa, HEC1 and SW620 cells in response to each drug. When combined with BOD1L1 deficiency, the lower dose of each drug caused a substantial further reduction in cell proliferation in U2OS cells, while HeLa cells were less affected and SW620 cells were not affected by the combination of BOD1L1 knockdown and taxol or Aurora kinase A inhibitor (Figure 8C, 8D, S5G). In line with the differential toxicity that we observed, U2OS cells with BOD1L1 knockdown formed micronuclei at a higher rate compared to control cells, while in SW620 cells there was no difference in micronuclei formation between control and BOD1L1 knockdown conditions (Figure S5F). Thus, the relative impact of BOD1L1 gene alteration on patient cancer survival is variable which is mirrored by a variable impact on proliferation of various cancer cell types following BOD1L1 depletion.

Finally, we tested the effect of BOD1L1 knockdown on the growth and sensitivity to taxol and Aurora kinase A inhibitor in non-transformed, diploid RPE1 cells. Immunoblots verify that the knockdown of BOD1L1 with each shRNA sequence was highly efficient (Figure S6A). Nevertheless, the knockdown of BOD1L1 did not sensitize RPE1 cells to taxol or the Aurora kinase A inhibitor at the dosages used. Indeed, RPE1 cells were more sensitive to taxol treatment than to the knockdown of BOD1L1, and they were not sensitive to the Aurora A kinase inhibitor (Figure S6).

## Discussion

Using an unbiased proteomic screen, we have discovered a pathway that regulates mitotic fidelity by controlling phosphatase activity directed at kinetochore substrates. In an unperturbed mitosis, BOD1L1 binds and inhibits PP2A-B56, helping to maintain moderate levels of phosphorylation of kinetochore substrates, thus decreasing k-MT attachment stability and ultimately supporting mitotic progression and high chromosome segregation fidelity. We envision a scenario that when CIN cancer cells are treated for prolonged periods with UMK57 to destabilize k-MT attachments, the adaptation response involves an increase in recruitment of Aurora kinase A to spindles through TPX2 which increases phosphorylation of BOD1L1 and relieves the negative regulation of PP2A activity. Relief of this constraint on PP2A leads to reduced phosphorylation of numerous kinetochore substrates at the k-MT interface which restores hyperstable k-MT stability and resumes the CIN phenotype (Figure 8E). This mechanism for adaptation to UMK57 is consistent with data showing that small changes in kinetochore phosphorylation can cause large changes in chromosome segregation fidelity^26–28^, and that the mechanism of adaptation is independent of the continued functional impact of UMK57 to activate MCAK^11^. Interestingly, it has been shown recently that cells adapt to oncogenic RAS/MAPK signaling by becoming increasingly dependent on BubR1/PP2A. This is because signaling through RAS/MAPK results in hyperactivity of mitotic kinases, reducing k-MT stability and increasing dependence on BubR1/PP2A to maintain viable levels of phosphorylation^29^. Thus, deregulation of phosphatase activity toward kinetochores substrates appears to be a common cellular response to pathways that change mitotic fidelity.

In support of our observations, the gene *cell proliferation regulating inhibitor of protein phosphatase 2A (CIP2A)* is fourth on the list of *BOD1L1* co-dependent genes from the CRISPR screens displayed in the Cancer Dependency Map Project^30,31^. CIP2A is a functional oncogene that inhibits the PP2A complex in various contexts including the DNA damage response, mitotic entry and mitotic progression by replacing the structural subunit of the PP2A complex, blocking phosphatase function^32,33^. Thus, the co-dependency between the two genes suggests that cell lines that are dependent on BOD1L1 for regulating phosphatase activity do not tolerate the loss of another gene that negatively regulates PP2A.

Notably, despite sharing a conserved domain, BOD1L1 and Bod1 do not bind PP2A through the same region. Our data show that BOD1L1 binds to PP2A with the middle and c-terminus of the protein which are not conserved in Bod1 (Figure 6D). Therefore, although the PP2A regulating function of Bod1 and BOD1L1 is conserved^22^, the mechanism of their interaction with PP2A is not. In addition, BOD1L1 has gained the ability to regulate DNA repair through binding Rif1 at the n-terminus^17^ and shifted the PP2A binding region to a large middle region of the protein plus the c-terminus. This genetic separation of DNA damage and PP2A regulating functions might explain the long c-terminal extension present in BOD1L1 compared to Bod1 and Bod1L2. Importantly, the phosphoacceptor sites on BOD1L1 that increase phosphorylation during the adaptation response to UMK57 also reside in the c-terminal domain of the protein and are not conserved with Bod1. Thus, the regulation of BOD1L1 by Aurora kinase A phosphorylation is distinct from processes that regulate Bod1.

A prevailing model differentiating the functional roles of Aurora A and B kinases in mitosis is that although they both use the same consensus motif to recognize substrates, their impact is different because they are spatially separated, with Aurora kinase A residing at spindle poles and on microtubules, and Aurora kinase B at centromeres. Although compelling, this model continues to be refined based on recent findings that some or perhaps even most non-CDK phosphorylation sites on Hec1 at kinetochores are phosphorylated by Aurora kinase A rather than Aurora kinase B^27,34,35^ and other data indicating that Aurora kinase B can fulfill its mitotic function without being targeted to centromeres in some systems^36^. Compared to Aurora kinase B, relatively little is known about how Aurora kinase A regulates k-MT attachments, although recent studies indicate that Aurora kinase A might also localize to kinetochores and phosphorylate substrates there or regulate kinetochore substrates on chromosomes proximal to the spindle pole^34,35^. Our data demonstrates that Aurora kinase A also regulates kinetochore substrate phosphorylation indirectly through regulation of BOD1L1 as a negative regulator of PP2A, and that this role for Aurora kinase A is responsible for changes in k-MT attachment stability induced by prolonged increase in MCAK activity caused by UMK57 treatment. In this context, an intriguing possibility emerges whereby the Aurora kinase A – BOD1L1 – PP2A axis identified in this work is activated by hypostable k-MT attachments (or hypostable spindle MTs more broadly) and induces a compensating increase in k-MT stability. Conversely, Aurora kinase B has a well-recognized role in decreasing k-MT stability. Together, overall k-MT stability (and perhaps total spindle MT stability and/or density) would be established through a lower bound regulated by Aurora kinase A activity and an upper bound regulated by Aurora kinase B activity. In support of this idea, TPX2 may serve as a sensor of MT stability to marshal Aurora kinase A as needed because TPX2 has been shown to sense and influence the conformational state of microtubules^37^ while promoting MT assembly by suppressing tubulin subunit off-rates^38^. Moreover, this role for Aurora kinase A in stabilizing spindle microtubules through BOD1L1 is consistent with many tumor cells overexpressing or having hyperactive Aurora kinase A^39–41^ and also being CIN as a consequence of hyperstable k-MT attachments. Our observations suggest that depending on the context, Aurora kinase A may destabilize k-MT attachments through direct kinetochore substrate phosphorylation or stabilize k-MT attachments through indirect regulation of PP2A activity.

Importantly, we found that the gene encoding BOD1L1 is occasionally altered in human cancers. Most of these alterations are point mutations, but some amplifications and deletions also occur (Figure 8A). In uterine endometrial cancer, the cancer that most commonly has BOD1L1 alterations, the alterations are associated with a reduced likelihood of death. The functional consequence of these mutations remains unknown, although it is likely that at least some of them cause a loss of BOD1L1 function. Thus, a reduced likelihood of death would be consistent with our findings that BOD1L1 knockdown results in a loss of fitness of various tumor lines and vulnerability to taxol or Aurora kinase A inhibitor (Figure 8C, 8D), which is corroborated by the mitotic arrest and occasional mitotic catastrophe observed in cells with BOD1L1 knockdown even in the absence of these drugs (Figure 5E, S3A). Since BOD1L1 also plays an important role in the cellular response to DNA replication stress and DNA repair^16,17^, these alterations may also contribute to the loss of cell fitness through that role. In contrast to patients with uterine endometrial cancer, alterations in BOD1L1 were associated with an increased risk of death in lung adenocarcinoma and in skin cutaneous skin melanoma. The clinically differing impacts of BOD1L1 alteration might depend on whether it is the mitotic functions and/or DNA repair functions that have been altered relative to the alterations at the root of a specific type of cancer. For example, both lung adenocarcinoma (if caused by smoking) and skin melanoma are driven by DNA damage caused by chemicals or UV light, thus possibly highlighting the role of BOD1L1 in DNA repair. Whereas in endometrial cancer, the cyclic high proliferation rate of this tissue in fertile women might highlight the role of BOD1L1 in maintaining accurate mitosis and avoiding CIN.

In summary, this work defines a new regulatory pathway in cells including Aurora kinase A, BOD1L1 and PP2A that is required for timely mitotic progression, faithful chromosome segregation, and regulation of proper microtubule stability. Our data expand on the previously known role for BOD1L1 as well as the previously known role of Aurora kinase A and provide insight into cellular responses that lead to adaptation to a small molecule suppressor of the CIN phenotype.

## Methods

### Cell Culture

SW620, HCT116, HeLa and U20S cells were obtained from the ATCC. HEC1 cells were a gift from Tyler Curiel. All cells were grown at 37°C in a humidified environment with 5% CO_2_. SW620, HeLa and U2OS cell lines were grown in Dulbecco’s modified Eagle medium (Corning® #15-017-CM) containing 10% FCS (Hyclone #82013-586), 250μg/L Amphotericin B (VWR #82026-728), 50 U/mL penicillin and 50 μg/mL streptomycin (ThermoFisher Scientific #15140122). HCT116 and HEC1 cells were grown in McCoy’s 5A medium with the same supplements. Cells were verified to be free of mycoplasma by frequent staining of plated cells with DAPI. No contamination was observed.

### Inhibitors and reagents

The CENP-E inhibitor GSK-923295 (MedChemExpress LLC # HY-10299) was used at 200 nM. NOC (VWR #80058-500) was used at 100-500 ng/ml. STLC (Tocris #2191) was used at 25 μM. ProTAME (Concept Life Sciences custom synthesis) was used at 25 μM. Okadaic acid (LC Labs O-5857) was used at 200 nM. RO-3306 (Selleck #S7747) was used at 10 μM. AZ3146 (R&D Systems #3994/10) was used at 2μM. ΒΙ2536 (Synthesized in-house) was used at 100 nM. MLN8237 (Selleck #S1133) was used at 10-50 nM. ZM447439 (Tocris Bioscience #2458/10) was used at 2 μM. UMK57 (Aobious #AOB8668) was used at 100 nM.

### Plasmids and cloning

The pLKO.1 – TRC cloning vector was a gift from David Root (Addgene plasmid # 10878; http://n2t.net/addgene:10878 ; RRID:Addgene_10878). The shRNA sequences were cloned into the AgeI and EcoRI sites of the plasmid using standard cloning techniques. The DNA constructs were then verified by Sanger sequencing and amplified by maxiprep (Qiagen #12162). Lentiviruses were produced by co-transfection of 293T/17 cells with pVSV-G and pdVPR29.1 plasmids. 48 hours later, the virus containing supernatant was harvested, briefly centrifuged to remove cell debris and frozen at -80°C until further use.

### Proteomic Screens

For the total protein and phospho-proteomic screen, ∼80% confluent cells were re-plated at a 1:10 dilution on 15 cm plates. 3 dishes were treated with DMSO and 3 with UMK57. 72 hours later, 2 dishes each of control and UMK57 treated cells were treated with 2.5 mM thymidine, and continuing DMSO/UMK57 treatment. 24 hours following thymidine addition, it was washed out with 2 washes of 5 ml PBS, and then fresh media added containing 100 ng/ml nocodazole, still containing DMSO/UMK57. 16 hours later the DMSO treated cells were treated for 1 hour with UMK57. The mitotic cells were harvested by shakeoff, centrifugation followed by a wash with 5 ml PBS, re-centrifugation and finally frozen at -80°C before mass spectrometry analysis. Separately, following the 72 hours DMSO/UMK57 treatment, the remaining dishes were trypsinized and re-plated on glass coverslips for microscopy analysis.

Sample preparation: Samples were prepared essentially as previously described^42,43^, with a few modifications. Cells were lysed in 8M urea, 200 mM EPPS, pH 8.5, supplemented with 1X Pierce protease and phosphatase inhibitors. The resuspended samples were passed through a 1.5-inch 21-gauge needle to ensure DNA were sheared. Protein concentration was determined for each sample with BCA. Samples were reduced with 5 mM TCEP at room temperature for 20 min, alkylated with 10 mM iodoacetamide at room temperature for 20 min in the dark, and then quenched with 10 mM DTT at room temperature for 20 min in the dark. Methanol-chloroform precipitation was performed prior to protease digestion. In brief, 4-parts neat methanol were added to each sample and vortexed, 1-part chloroform was added to the sample and vortexed, and 3-parts water was added to the sample and vortexed. The sample was centrifuged at 14,000 RPM for 2 min at room temperature and subsequently washed once with 100% methanol. Samples were resuspended in 200 mM EPPS, pH 8.5 and digested at room temperature for 14 h with Lys-C protease at a 50:1 protein-to-protease ratio. Trypsin was then added at a 100:1 protein-to-protease ratio and the reaction was incubated for 6 h at 37°C.

Fe^3^+-NTA phosphopeptide enrichment: Following digestion with Lys-C and trypsin, peptides were desalted by using 100mg SepPak columns. The elutions were dried via vacuum centrifugation and the phosphopeptides were enriched with the High-Select Fe3+-NTA Phosphopeptide Enrichment Kit according to manufacturer’s specifications using approximately 4 mg protein digest per enrichment column. The elutions were dried via vacuum centrifugation, while the flow-throughs were saved for subsequent whole proteome analysis.

TMT labeling: In general, we estimate that at most 20 ug of phosphopeptides were enriched from 4 mg of total peptide. We added 40 ug of TMT 6-plex reagent (Thermo-Fisher) to the peptides along with acetonitrile to again achieve a final acetonitrile concentration of approximately 30% (v/v) in a total volume of 50 µL. Following incubation at room temperature for 1 h, the reaction was quenched with hydroxylamine to a final concentration of 0.3% (v/v). The sample was vacuum centrifuged to near dryness and subjected to C18 solid-phase extraction (SPE, Sep-Pak). Off-line basic pH reversed-phase (BPRP) fractionation: We fractionated the peptide samples using BPRP HPLC. We used an Agilent 1200 pump equipped with a degasser and a UV detector. Peptides were subjected to a 50-min linear gradient from 5% to 35% acetonitrile in 10 mM ammonium bicarbonate pH 8 at a flow rate of 0.6 mL/min over an Agilent 300Extend C18 column (3.5 μm particles, 4.6 mm ID and 250 mm in length). The peptide mixture was fractionated into a total of 96 fractions, which were consolidated into 12 super-fractions.

Samples were subsequently acidified with 1% formic acid and vacuum centrifuged to near dryness. Each consolidated fraction was desalted by StageTip, and reconstituted in 5% acetonitrile, 5% formic acid for LC-MS/MS processing.

Liquid chromatography and tandem mass spectrometry: Mass spectrometric data were collected on an Orbitrap Lumos mass spectrometer coupled to a Proxeon NanoLC-1200 UHPLC (ThermoFisher Scientific). The 100 µm capillary column was packed in-house with 35 cm of Accucore 150 resin (2.6 μm, 150Å; ThermoFisher Scientific). Data were acquired for 180 min per fraction. MS1 spectra were acquired at 120 K resolving power for a maximum of 50 ms in the Orbitrap, and features were filtered for monoisotopic peak assignment and charge states 2-6. MS2 spectra were acquired by selecting the top 10 most abundant features. MS2 spectra were acquired via collisional induced dissociation (CID), in the ion trap with an automatic gain control (AGC) of 20K, quadrupole isolation width of 0.5 m/z and a maximum ion time of 120 ms with Multistage Activation turned on. For MS3 acquisition, a synchronous precursor selection (SPS) of 10 fragments ions was acquired for a maximum of 150 ms with an AGC of 300K and a normalized collision energy of 65.

Data analysis: Database searching included all entries from the mouse UniProt Database (downloaded: 2014). The database was concatenated with one composed of all protein sequences for that database in the reversed order^44^. Raw files were converted to mzXML, and monoisotopic peaks were re-assigned using Monocle^45^. Searches were performed with SEQUEST^46^ using a 20-ppm precursor ion tolerance. TMT labels on lysine residues and peptide N-termini (+229.1629 Da), as well as carbamidomethylation of cysteine residues (+57.021 Da) were set as static modifications, while oxidation of methionine residues (+15.995 Da) and phosphorylation (+79.966) were set as variable modifications. Peptide-spectrum matches (PSMs) were adjusted to a 1% false discovery rate (FDR) using a linear discriminant after which proteins were assembled further to a final protein-level FDR of 1% analysis^47^. Ascore was used to determine site localization^48^, with a score of 13 denoting 95% confidence for a specified phosphorylation site. Phosphorylation sites were quantified by summing reporter ion counts across all matching PSMs. Peptides were filtered to include only those with a summed signal-to-noise (SN) ≥ 100 across all TMT channels as well as an isolation specificity (“isolation purity”) of at least 0.5.

### Whole Proteome Profiling

Sample preparation for whole proteome profiling: Flow-throughs from the phospho-enrichment described above, 50 µg per replicate, were used for the whole proteome work for each sample. TMT labeling: 120 µg of TMTpro reagents (Thermo-Fisher; Lot # WL338745) were added to the peptides (50 µg) along with acetonitrile to achieve a final acetonitrile concentration of approximately 30% (v/v) in a total volume of 100 µL. Following incubation at room temperature for 1 h, the reaction was quenched with hydroxylamine to a final concentration of 0.3% (v/v). The sample was vacuum centrifuged to near dryness and subjected to C18 solid-phase extraction (SPE, Sep-Pak).

Off-line basic pH reversed-phase (BPRP) fractionation: We fractionated the peptide samples using BPRP HPLC. We used an Agilent 1200 pump equipped with a degasser and a UV detector. Peptides were subjected to a 50-min linear gradient from 5% to 35% acetonitrile in 10 mM ammonium bicarbonate pH 8 at a flow rate of 0.6 mL/min over an Agilent 300Extend C18 column (3.5 μm particles, 4.6 mm ID and 250 mm in length). The peptide mixture was fractionated into a total of 96 fractions, which were consolidated into 24 super-fractions (in a checkerboard-like pattern) of which 12 were used for data acquisition. Samples were subsequently acidified with 1% formic acid and vacuum centrifuged to near dryness. Each consolidated fraction was desalted by StageTip, and reconstituted in 5% acetonitrile, 5% formic acid for LC-MS/MS processing.

Liquid chromatography and tandem mass spectrometry. Mass spectrometric data were collected on an Orbitrap Fusion mass spectrometer coupled to a Proxeon NanoLC-1200 UHPLC (ThermoFisher Scientific). A 100 µm capillary column was packed in-house with 35 cm of Accucore 150 resin (2.6 μm, 150Å; ThermoFisher Scientific). Data were acquired for 180 min per fraction. MS1 spectra were acquired at 120 K resolving power for a maximum of 100 ms in the Orbitrap, and features were filtered for monoisotopic peak assignment and charge state of 2. MS2 spectra were acquired by selecting the top 10 most abundant features. MS2 spectra were acquired via collisional induced dissociation (CID), in the ion trap with an automatic gain control (AGC) of 9K, quadrupole isolation width of 0.7 m/z at Rapid Scan Rate. For MS3 acquisition, a synchronous precursor selection (SPS) of 10 fragments ions was acquired for a maximum of 200 ms with an AGC of 200K and a normalized collision energy of 55.

Data analysis. Searches were performed as described above but without phosphorylation variable modifications. Proteins were quantified by summing reporter ion counts across all matching PSMs. More specifically, reporter ion intensities were adjusted to correct for the isotopic impurities of the different TMTpro reagents according to manufacturer specifications. Peptides were filtered to include only those with a summed signal-to-noise (SN) ≥ 100 across all TMT channels and isolation specificity of at least 0.5. The signal-to-noise (S/N) measurements of peptides assigned to each protein were summed (for a given protein) and each protein abundance measurement was scaled, such that the summed signal-to-noise for that protein across all channels equals 100, thereby generating a relative abundance (RA) measurement.

### Transfections

200,000 SW620 cells were plated on glass coverslips in 12-well plates. 24 hours following plating, they were transfected with 1 μg plasmid DNA and 2.5 μl Lipofectamine^TM^ 2000 (ThermoFisher #11668027) in a total of 1 ml Opti-MEM^TM^ media (ThermoFisher #31985070). Four hours following transfection, the media was changed for DMEM containing 2μg/ml puromycin. The cells were then assayed 48-72 hours following addition of puromycin.

### SDS-PAGE and Western blot

Gels and blots were performed as described previously ^49^. Prior to loading samples on gel, 4X Laemmli buffer (200 mM Tris pH 6.8, 4% SDS, 40% glycerol, 4% 2-mercaptoethanol, 0.12 mg/ml bromophenol blue) was added to a final dilution of 1X, and the samples were boiled for 5 minutes. Proteins were separated on SDS-PAGE gel using stacking gel (4% 29:1 acrylamide: Bis-acrylamide, 125mM Tris pH 6.8, 0.1% SDS, 0.1% ammonium persulfate, 0.1% TEMED) and (8-15% 29:1 acrylamide: Bis-acrylamide, 400 mM Tris pH 8.8, 0.1% SDS, 0.1% ammonium persulfate, 0.1% TEMED) at 120V until the bromophenol blue has run off or longer, as needed. Transfer onto nitrocellulose membrane was performed for at least 24 hours at 30V under wet conditions in 1X transfer buffer (14.4 g/L glycine, 3.0 g/L Tris, 20% methanol). Conditions for western blots includes the use of 5% nonfat dry milk in TBS-T 0.5% (50 mM Tris [pH 7.2], 150 mM NaCl, 0.5% Tween 20) for blocking and TBS-T 0.5% for washing. 4 washes for 10 minutes each were performed after primary and secondary antibody incubation periods. The bands were visualized by enhanced chemiluminescence using Clarity (Bio-Rad #1705060) using a Bio-Rad ChemiDoc MP.

### Immunoprecipitation

Protein-G beads were first crosslinked to the antibody by incubating the beads and antibody overnight on a rotating platform. The beads were then washed with PBS and then crosslinking solution (20mM dimethyl pimelimidate, 0.3M HEPES in ice cold water). The beads were then incubated in crosslink solution for 10 minutes at room temperature. The solution is then discarded, and the incubation repeated in fresh crosslink buffer, which is performed three times. The beads are then washed in PBS twice and twice with lysis buffer prior to use.

SW620 cells from confluent 15 cm dishes were split 1:5. 48 hours later, 2.5 mM thymidine was added. 24 hours following thymidine addition, it was washed out with 2 5ml washes with PBS. Fresh media containing 100 ng/ml nocodazole was then added. 16 hours later, the cells were harvested by scraping, washed once with cold PBS, and lysed in the following buffer: 0.5% NP40, 100 mM NaCl, 50 mM tris pH 7.4, 10 mM glycerol 2-phosphate. Cells were lysed on ice for 20 minutes before centrifuging at 15,000 x g for 20 minutes at 4°C. The supernatant was transferred to a fresh tube, and input sample set aside. 10 μl of packed volume protein G beads cross-linked to antibody were added to each sample for 2 hours on a rotating platform at 4°C. The beads were then washed 5 times in lysis buffer and dried using a 30 g needle attached to a vacuum line. 30-100 μl of 1X sample buffer were added to the beads before elution by boiling and loading on gel.

### Identification of BOD1L1 interacting proteins

Excised gel bands were cut into approximately 1 mm^3^ pieces. Gel pieces were then subjected to a modified in-gel trypsin digestion procedure^50^. Gel pieces were washed and dehydrated with acetonitrile for 10 min. followed by removal of acetonitrile. Pieces were then completely dried in a speed-vac. Rehydration of the gel pieces was with 50 mM ammonium bicarbonate solution containing 12.5 ng/µl modified sequencing-grade trypsin (Promega, Madison, WI) at 4°C. After 45 min., the excess trypsin solution was removed and replaced with 50 mM ammonium bicarbonate solution to just cover the gel pieces. Samples were then placed in a 37°C room overnight. Peptides were later extracted by removing the ammonium bicarbonate solution, followed by one wash with a solution containing 50% acetonitrile and 1% formic acid. The extracts were then dried in a speed-vac (∼1 hr). The samples were then stored at 4°C until analysis. On the day of analysis the samples were reconstituted in 5 – 10 µl of HPLC solvent A (2.5% acetonitrile, 0.1% formic acid). A nano-scale reverse-phase HPLC capillary column was created by packing 2.6 µm C18 spherical silica beads into a fused silica capillary (100 µm inner diameter x ∼30 cm length) with a flame-drawn tip^51^. After equilibrating the column each sample was loaded via a Famos auto sampler (LC Packings, San Francisco CA) onto the column. A gradient was formed and peptides were eluted with increasing concentrations of solvent B (97.5% acetonitrile, 0.1% formic acid). As peptides eluted they were subjected to electrospray ionization and then entered into an LTQ Orbitrap Velos Pro ion-trap mass spectrometer (Thermo Fisher Scientific, Waltham, MA). Peptides were detected, isolated, and fragmented to produce a tandem mass spectrum of specific fragment ions for each peptide. Peptide sequences (and hence protein identity) were determined by matching protein databases with the acquired fragmentation pattern by the software program, Sequest (Thermo Fisher Scientific, Waltham, MA)^46^. All databases include a reversed version of all the sequences and the data was filtered to between a one and two percent peptide false discovery rate.

### GST pulldowns

We performed GST-pulldowns essentially as described previously^17^. Briefly, SW620 mitotic nuclear extracts were incubated with 1 μg of GST-fusion protein or purified GST as a control on a rotating platform for 3 hours. Bound proteins were then purified via binding to glutathione Sepharose (GE Healthcare) and washed. Samples were then analyzed by Western blot.

### RNA interference

Gene knockdown was performed by cloning the following shRNA sequences into the PLKO1 vector and then transfecting cells with 1μg of plasmid DNA and 2.5μl lipofectamine 2000 per ml according to the manufacturer’s directions. Media was changed after 5 hours, and puromycin added after 24 hours.

BOD1L1-1: 5’- GCCAATGATGCCATGTCGATA-3’

BOD1L1-2: 5’- ACTCGCATGTATCCAAGTAAA-3’

### Immunofluorescence

Following treatments, cells were pre-extracted in PHEM buffer (60 mM PIPES, 25 mM HEPES, pH 6.9, 10 mM EGTA, and 4 mM MgSO_4_, 1% Triton X-100, 10 mM glycerol 2-phosphate) for 10 minutes at 4°C and then fixed in 4% paraformaldehyde in PBs at room temperature for 20 minutes. The remaining steps were performed at room temperature. The cells were then washed twice with PBS for before blocking with ADB (10% serum, 0.1% Triton X-100 in PBS) for 15 minutes. Primary antibodies were then diluted in ADB and used at the indicated dilution. Secondary antibodies were diluted in ADB. After incubation with antibodies, the cells were washed three times with PBS 0.1% Triton X-100. Finally, the cells were counterstained with DAPI and mounted on slides using ProLong^TM^ Gold antifade reagent (ThermoFisher Scientific #P36934).

### Microscopy

Images were acquired with a Nikon Eclipse Ti microscope equipped with a cooled charge-coupled device Clara camera (Andor Technology) controlled by Nikon NIS-Elements software version 4.30.02. Images were acquired in 0.15-0.4 μm sections using a plan apo 1.4 numerical aperture 100X (kinetochore analysis) or 60X (lagging chromosome analysis) oil-immersion objective using 1X1 binning. Samples were illuminated using an X-cite light source (Excelitas Technologies Corp). All image analysis, adjustment and cropping was performed using Fiji software ^52^. Image deconvolution was performed using Nikon Elements batch deconvolution software version 5.21.00 on automatic mode. All kinetochore intensity analysis was performed on the raw images, except for microtubule intensity determination for which deconvolution was performed prior to analysis. Line scans were performed on the deconvoluted images in Fiji and the data was then exported to Graphpad Prism. All images of BOD1L1 selected for presentation are deconvolved single optical slices. Images of other stained proteins are deconvolved maximum intensity projections. Where shown, insets are from single optical Z-slices. Images were selected to represent the mean quantified data. Quantification of kinetochore staining intensity was performed by first determining the brightest plane for which a given kinetochore appears. Then, an outline was drawn using the ellipse tool. The integrated intensity was determined by multiplying the average intensity by the area of the kinetochore. Background levels of staining were determined by saturating the brightness and contrast settings to find the darkest spot within the chromatin containing area and were subtracted from the intensity of the kinetochore. Aurora kinase A pT288 was quantified by first creating a sum projection of the image. Then, an outline was drawn around the whole cell, and the fluorescence intensity measured. The background-subtracted, normalized integrated intensities for all cells were then plotted. To quantify levels of cold-stable microtubules, images were first deconvolved. Then, A sum-intensity projection was created in Fiji. Viewing the ACA channel, a rectangle was drawn that was 2 μm wide in the pole-pole axis, and as long as the furthest centromeres in the metaphase plate. Then, switching to the tubulin channel, the average tubulin intensity was measured in the rectangle. From the average intensity value, the minimum intensity value was used as the background level and subtracted. The Background-subtracted intensity values were then averaged across all of the cells that were analyzed for each condition and experimental repeat.

For live-cell microscopy, cells were transfected with 0.5 μg of mCherry-H2B-expressing plasmid together with pLKO vectors. After selection with puromycin, the cells were washed twice with PBS, given fresh medium and moved to a Tokai Hit incubation chamber with CO_2_ and temperature control on the microscope. The cells were imaged every 5 minutes at 20X magnification for 48 hours with 2X2 binning.

### Antibodies

The following antibodies were used for immunofluorescence (IF) and/or immunoblotting (IB): human anti-ACA (Geisel School of Medicine; IF at 1:2000), mouse anti-Hec1 (Santa Cruz C-11; IF,IB at 1:1000), mouse anti-PP2AR1A (Santa Cruz SC-74580; IB at 1:500), mouse anti-PP2AC (Santa Cruz SC-166034; IB at 1:500), rabbit anti-PP2A B56 Alpha (MyBioSource.com MBS8524809; IB at 1:500), mouse anti-PP2A B56 Beta (Santa Cruz E-6; IB at 1:500), mouse anti-PP2A B56 Gamma (Santa Cruz A-11; IB at 1:500), mouse anti-PP2A B56 Delta (Santa Cruz H-11; IB at 1:500), rabbit anti-PP2A B56 Epsilon (Aviva Systems Biology RP56694_P050; IB at 1:500), rabbit anti-Hec1 pS44 (gift of Jennifer DeLuca; IF at 1:500)^53^, rabbit anti DSN1 pS100 (gift of Iain Cheeseman; IF at 1:250)^54^, rabbit anti-KNL1 pT943/pT1155 (Cell Signaling #40758), mouse anti-α-Tubulin DM1α (Sigma-Aldrich; IF at 1:4000-1:10,000), mouse anti-Aurora A (Cell Signaling Technology 1F8; IF at1:1000), Rabbit anti-Aurora A pT288 (Cell Signaling Technology C39D8; IF at 1:500), Rabbit anti-TACC3 (gift of Jordan Raff; IF at 1:1000)^55^, mouse anti-TPX2 (Cell Signaling Technology D9Y1V; IF at 1:1000), rabbit anti-Aurora B pT232 (Rockland 600-401-677; IF/IB at 1:1000), rabbit anti-MCAK pS95 (Abcam #AB74146; IF/IB at 1:500), rabbit anti-Hec1 pT31^26^ (IF at 1:100,000, WB at 1:1000), rabbit anti-BOD1L1 (IF at 1:1000) (gift of Grant Stewart^16^), rabbit anti-BOD1L1 (Genetex #GTX119946; WB at 1:500). Secondary antibodies used were highly cross-adsorbed Alexa Fluor® 488, 594 and 647 raised in donkey or goat (Invitrogen; IF at 1:1000), horseradish peroxidase (Bio-Rad; IB at 1:10000-10,0000), Mouse TrueBlot® ULTRA: rat Anti-Mouse IgG (Rockland 18-8817-33; IB at 1:1000), Anti-IgG (H+L) Goat Polyclonal Antibody (Horseradish Peroxidase), Peroxidase AffiniPure Goat Anti-Rabbit IgG (H+L) (IB at 1:5000), Goat Polyclonal Antibody (Horseradish Peroxidase), Peroxidase AffiniPure Goat Anti-Mouse IgG (H+L) (IB at 1:5000).

### Statistical analysis of cell biology data

All statistical tests were performed using Graphpad Prism version 8.4.2. Except where indicated, all experiments were performed as 2-3 independent biological repeats. In most IF experiments, 20 cells per repeat were analyzed, and 25 kinetochores for each cell measured. Experiments with other numbers are noted in the figure legends. For anaphase error rates, 100 anaphases per condition were assessed for each of 3 independent biological experiments. Statistical significance was calculated between indicated conditions using two-tailed *t*-tests or Dunnett’s multiple comparison test. For these tests, the data distribution was assumed to be normal but this was not formally tested.

### Study population data, and statistical analysis

The cBioPortal for Cancer Genomics was used to query and visualize *BOD1L1* alterations in data from The Cancer Genome Atlas (TCGA). Then, TCGA PanCancer Atlas cancer specific clinical and *BOD1L1* alteration data was downloaded via cBioPortal for Cancer Genomics for cancer types where *BOD1L1* alteration was most frequent: uterine corpus endometrial carcinoma TCGA-UCEC PCA (n= 529), lung adenocarcinoma TCGA-LUAD (n = 667), cutaneous melanoma TCGA-SKCM (n = 448), stomach adenocarcinoma TCGA-STAD (n = 440), and lung squamous cell carcinoma TCGA-LUSC (n = 487). All statistical analysis was conducted in R version 4.2.1. Survival analysis using Kaplan-Meier plots and Cox Proportional-Hazards models was conducted using R package *survival* and plotted using R package *survminer*. Univariate Kaplan-Meier survival strata were plotted for altered *BOD1L1* status in each cancer type to assess five-year overall survival status. Cox Proportional Hazards models for each cancer type tested association of *BOD1L1* alteration status with adjustment for patient age, sex (excepting uterine), and tumor stage or grade. Using Fisher’s exact tests and patient data from all TCGA-PanCancer Atlas cancer datasets (n = 10,967 from 32 cancer types), we tested whether *BOD1L1* alterations were exclusive of alterations to other Aurora A-BOD1L1-PP2A gene pathway members.

### Data availability

The cell biology datasets generated during and/or analysed during the current study are available from the corresponding author on reasonable request. The MS datasets will be uploaded to an appropriate public repository upon acceptance of the article.

## Acknowledgements

We thank Grant Stewart, Jennifer Delucca, Iain Cheeseman, Tyler Curiel, Jose Teodoro and Adrian Saurin for providing reagents. We also thank Sanjukta Guha Thakurta (Thermo Fisher Center for multiplexed proteomics, Harvard University) and Ross Tomaino (Taplin Mass Spectrometry Facility, Harvard University) for assistance with the proteomics analysis. We also thank the members of the Compton laboratory for helpful discussions and feedback. TJK was supported by fellowships from the Fonds de Recherche du Québec – Santé and the Canadian Institutes of Health Research. The laboratory of SB is supported by grants from the Canadian Institutes of Health Research (PJT-156193) and Natural Sciences and Engineering Research Council of Canada (NSERC; RGPIN-2017-04649). This work was also supported by grants from the National Institutes of Health #GM051542 to DAC. The authors acknowledge the following Shared Resources facilities (Bio-Rad ChemiDoc MP) at the Norris Cotton Cancer Center at Dartmouth with NCI Cancer Center Support Grant 5P30 CA023108-40.

## Author Contributions

Conceptualization: DAC and TJK. Investigation: TJK, IMV, MRH. Formal analysis: TJK, IMV. Methodology: TJK, DAC. Funding acquisition: TJK, DAC. Supervision: BCC, SB, DAC. Writing-original draft: TJK. Writing-review & editing: TJK, MRH, BCC, DAC. Project administration: DAC.

## Materials & Correspondence

Any requests should be addressed to DAC

---

**Figure S1:**
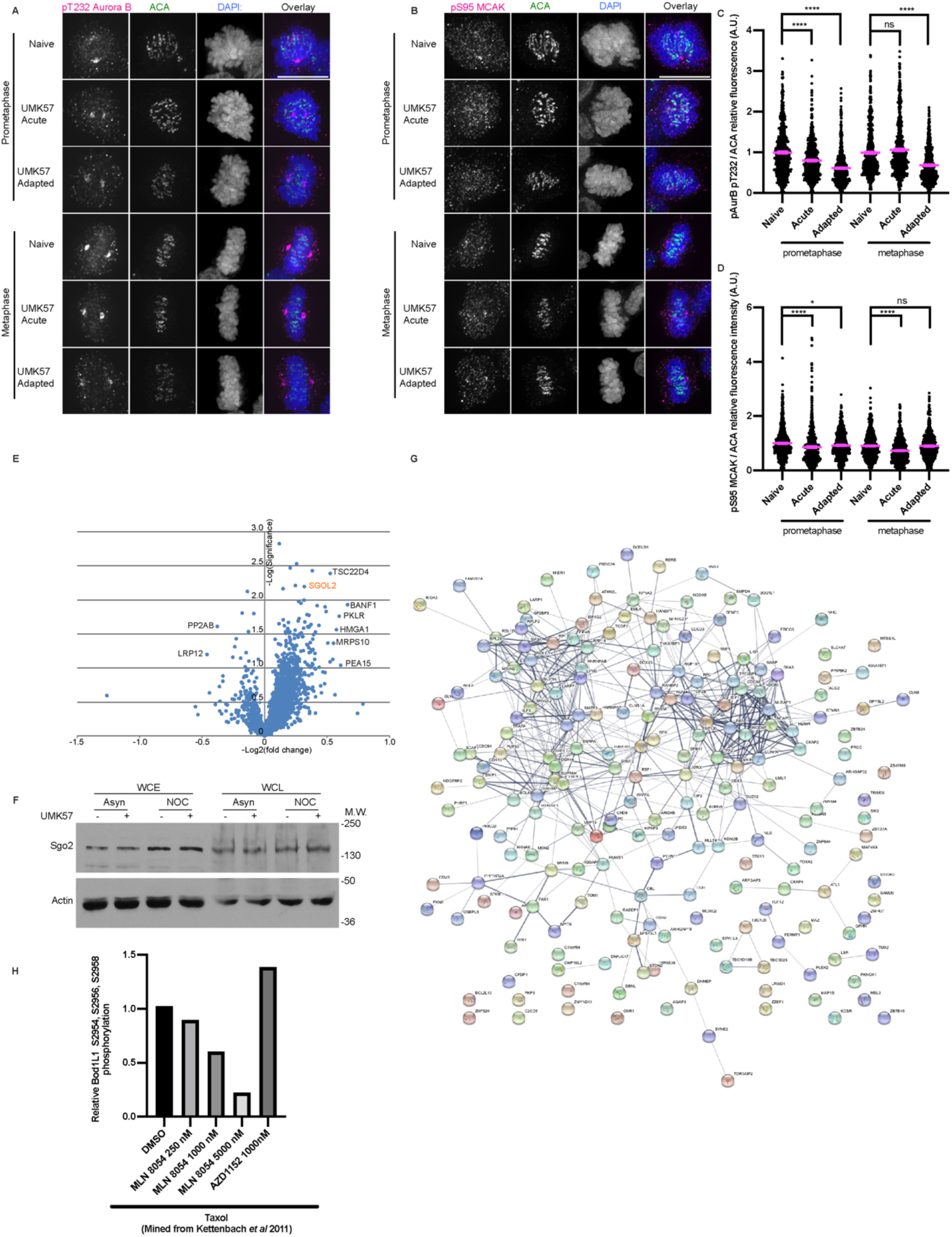
Changes in UMK57 adapted cells; Adaptation to UMK57 does not happen through changes to protein level. A) SW620 cells were first treated for 3 days with UMK57 or DMSO control. They were then re-plated on glass coverslips. After 24 hours, the cells were fixed and stained for Aurora B pT232, ACA and DAPI. A single experiment was performed. The scale bar is 10μm. B) SW620 cells were prepared as in (A) but stained for MCAK pS95 and ACA. A single experiment was performed. The scale bar is 10μm. C) Quantification of the relative kinetochore pT232 Aurora B/ACA intensities from the conditions from panel (A). The condition with the lowest level of pT232 Aurora B/ACA was set to 1 and the other conditions shown as fold-changes. 30 kinetochores were quantified from each of 20 cells from a single experiment. Error bars indicate the mean +/- SEM. Statistical significance was calculated between the indicated conditions using Dunnett’s multiple comparison test. D) As for C but quantifying the relative kinetochore pS95 MCAK/ACA intensities from panel (B). E) Volcano plot analysis of the total protein level proteomic screen results. Proteins of potential interest are highlighted in orange. Statistical significance was calculated using two-tailed *t*-tests. F) STRING analysis of the statistically significant (*p* < 0.1) top-100 positively changed phosphorylated proteins and top-100 negatively changed phospho-proteins showing gene association networks. G) SW620 cells were first treated for 3 days with UMK57 or DMSO control. They were then allowed to remain asynchronous or were synchronized by thymidine-nocodazole. The cells were then harvested and lysed. The proteins were separated by SDS-PAGE, transferred to nitrocellulose membrane and finally blotted as indicated. The shown blots are representative of 3 independent experiments.

**Figure S2:**
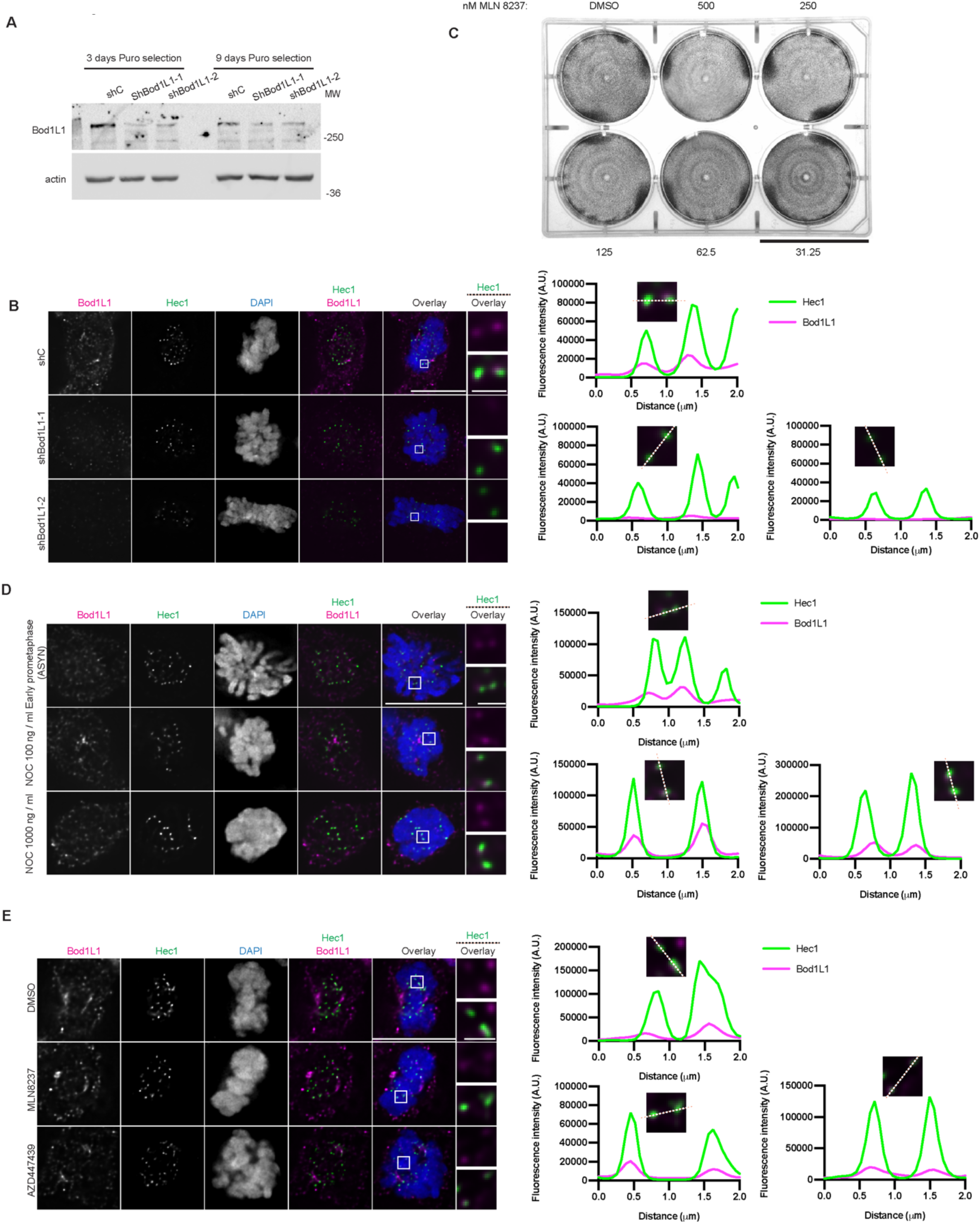
Validation of BOD1L1 antibody and shRNAs; BOD1L1 does not require Aurora A, Aurora B signaling or microtubules for kinetochore localization. A) SW620 cells were transfected with control shRNA or shRNA against BOD1L1 and selected with puromycin. The cells were then harvested at the indicated time points, lysed and the proteins separated by SDS-PAGE. They were then transferred to nitrocellulose membrane and blotted as indicated. B) Immunofluorescence images from an asynchronous population of SW620 transfected with control shRNA or shRNA against BOD1L1 and selected with puromycin. The cells were stained for BOD1L1, Hec1 and DAPI. The BOD1L1 and Hec1 channels were adjusted evenly for brightness and contrast for presentation. The DAPI channel of each condition was adjusted independently. Representative images from 2 independent biological experiments are shown. The scale bars of main images are 10 μm, and insets are 1 μm. C) SW620 cells were plated. 24 hours following plating, various concentrations of MLN8237 were added as indicated. Following 3 days of treatment, the cells were fixed and stained with crystal violet. Representative images from 2 independent experiments are shown. The scale bar is 4 cm. D) Immunofluorescence images from an asynchronous population of SW620 transfected with control shRNA or shRNA against BOD1L1 and selected with puromycin. The cells were overnight with the indicated concentrations of nocodazole. The cells were then fized and stained for BOD1L1, Hec1 and DAPI. The BOD1L1 and Hec1 channels were adjusted evenly for brightness and contrast for presentation. The DAPI channel of each condition was adjusted independently. Representative images from 2 independent biological experiments are shown. The scale bars of main images are 10 μm, and insets are 1 μm. E) Immunofluorescence images from an asynchronous population of SW620 transfected with control shRNA or shRNA against BOD1L1 and selected with puromycin. The cells were then treated with DMSO control or Aurora A or Aurora B inhibitors for 90 minutes prior to fixation. The cells were stained for BOD1L1, Hec1 and DAPI. The BOD1L1 and Hec1 channels were adjusted evenly for brightness and contrast for presentation. The DAPI channel of each condition was adjusted independently. Representative images from 2 independent biological experiments are shown. The scale bars of main images are 10 μm, and insets are 1 μm.

**Figure S3:**
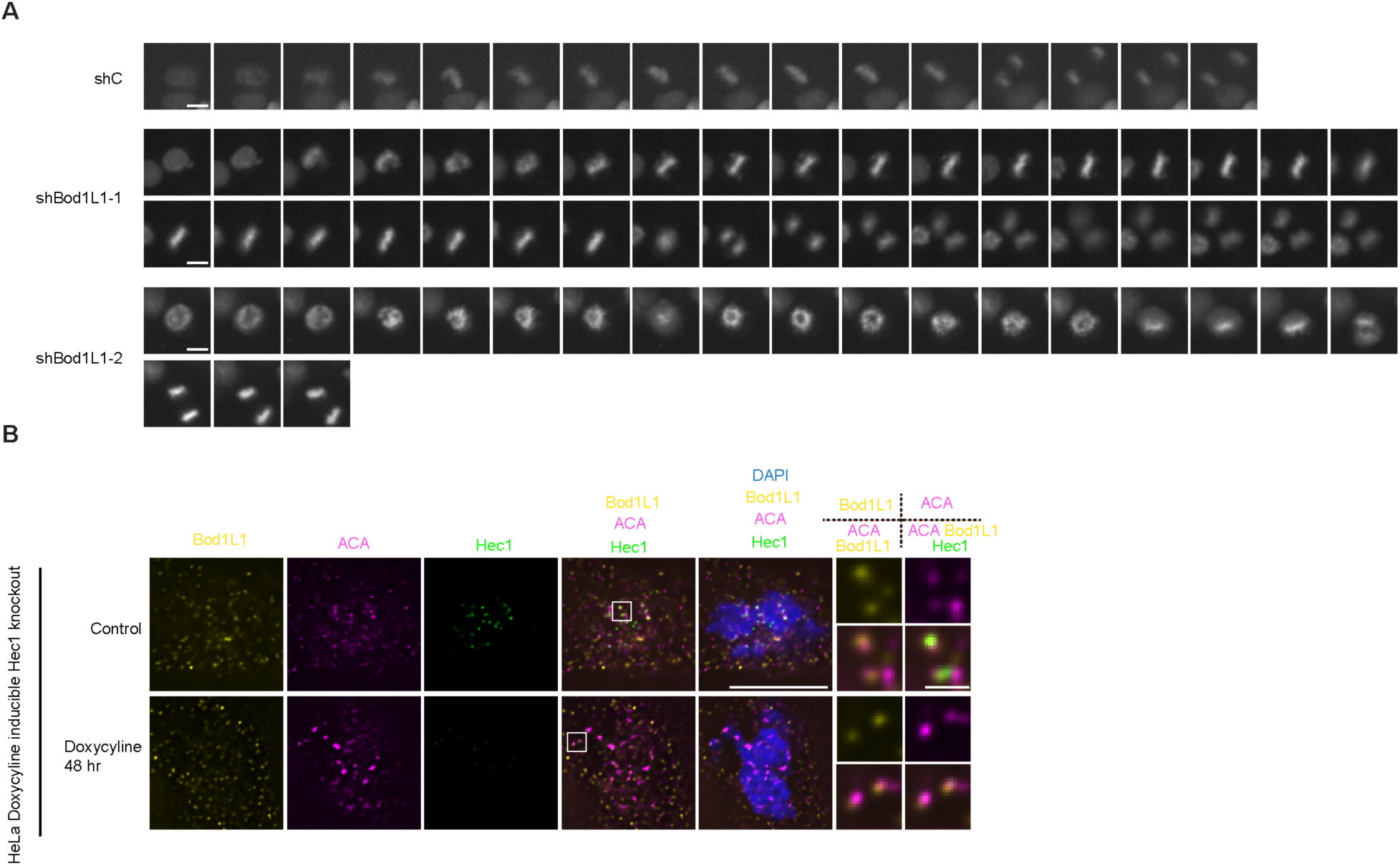
Still images of live-cell movies; Hec1 is not required for BOD1L1 localization to kinetochores. A) SW620 cells were co-transfected with control shRNA or shRNA against BOD1L1 together with mCherry-H2B and selected with puromycin. After 72 hours of selection, time lapse imaging was started at 5-minute intervals. Representative Images from 3 independent experiments are shown. The images were independently adjusted for brightness and contrast for presentation. The scale bars are 10 μm. B) HeLa cells were plated on glass coverslips. 24 hours after plating, doxycycline or water was added to the wells. 48 hours following addition of doxycycline, the cells were fixed and stained as indicated. Representative Images from 2 independent experiments are shown. The images were independently adjusted for brightness and contrast for presentation. The scale bars of main images are 10 μm, and insets are 1 μm.

**Figure S4:**
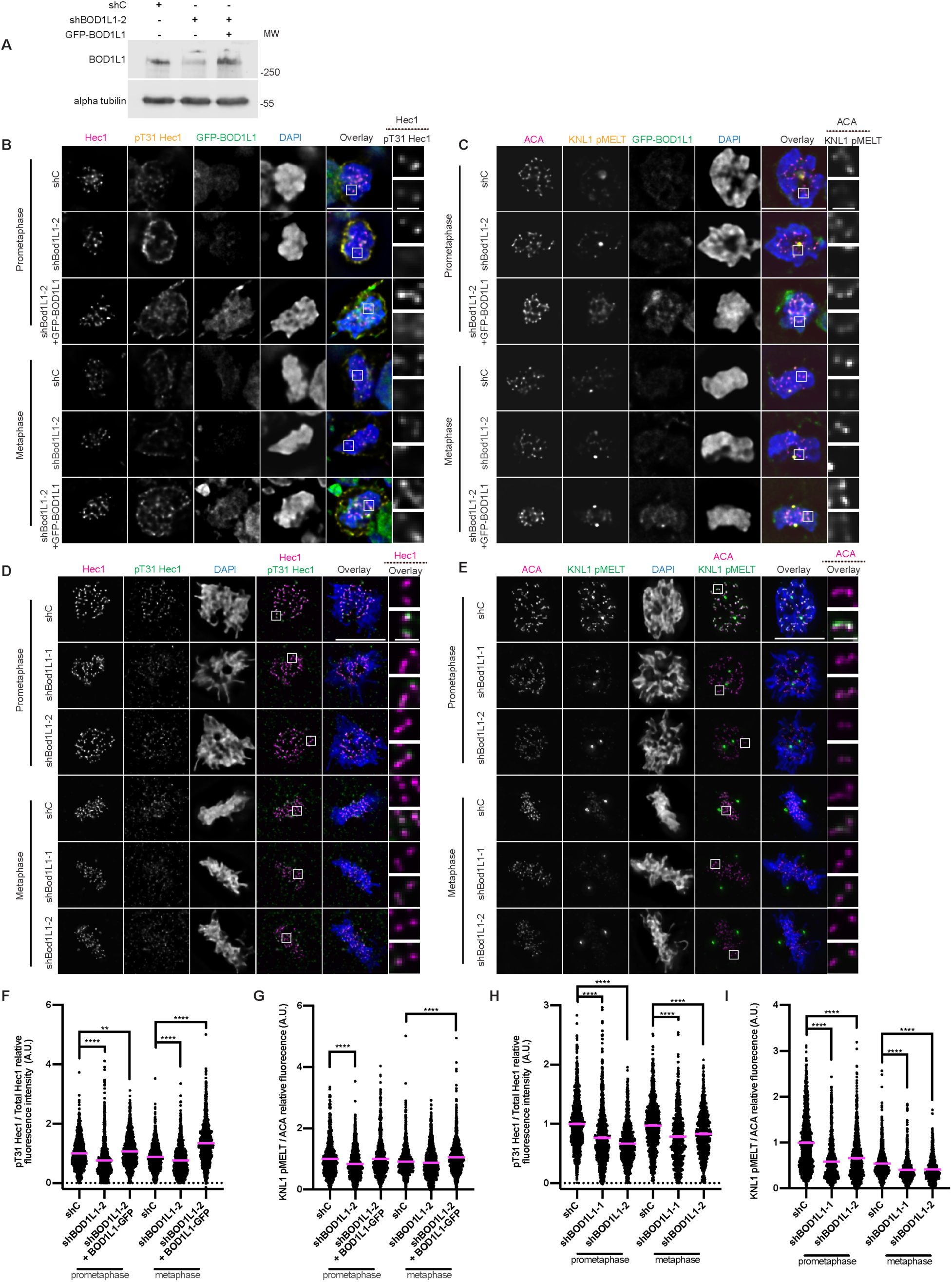
Overexpression of GFP-BOD1L1 rescues depletion of BOD1L1. A) SW620 cells were plated. They were then transfected as indicated. 24 hours following transfection, puromycin was added. The cells were harvested 72 hours after transfection and lysed. The proteins were separated by SDS-PAGE, transferred to nitrocellulose membrane and finally blotted as indicated. A single experiment was performed. B) SW620 cells were plated on glass coverslips. They were then transfected as indicated. 24 hours following transfection, puromycin was added. 72 hours following transfection, the cells were fixed and stained as indicated. Representative Images from 2 independent experiments are shown. The images were independently adjusted for brightness and contrast for presentation, except for DAPI, which was adjusted independently. The scale bars of main images are 10 μm, and insets are 1 μm. C) As for (B), but stained as indicated. D) RPE1 cells were plated on glass coverslips. They were then transduced with lentivirus as indicated. 24 hours following transduction, puromycin was added. 72 hours following transduction, the cells were fixed and stained as indicated. Representative Images from 2 independent experiments are shown. The images were independently adjusted for brightness and contrast for presentation, except for DAPI, which was adjusted independently. The scale bars of main images are 10 μm, and insets are 1 μm. E) As for D but stained as indicated F) Quantification of the relative kinetochore pT31 Hec1/Hec1 intensities from the conditions from panel (B). The condition with the lowest level of pT31 Hec1/Hec1 was set to 1 and the other conditions shown as fold-changes. 25 kinetochores were quantified from each of 20 cells for each of 2 independent biological repeats. Error bars indicate the mean +/- SEM. Statistical significance was calculated between the indicated conditions using Dunnett’s multiple comparison test G) Quantification of the relative kinetochore KNL1 pMELT intensities from panel (C) as for panel (F). H) Quantification of the relative pT31 Hec1/Hec1 intensities from the conditions from panel (D) as for panel (F). 25 kinetochores were quantified from at least 22 cells over two independent biological repeats. I) Quantification of the relative KNL1 pMELT intensities from panel (E) as for panel (F). 25 kinetochores were quantified from at least 27 cells over two independent biological repeats.

**Figure S5:**
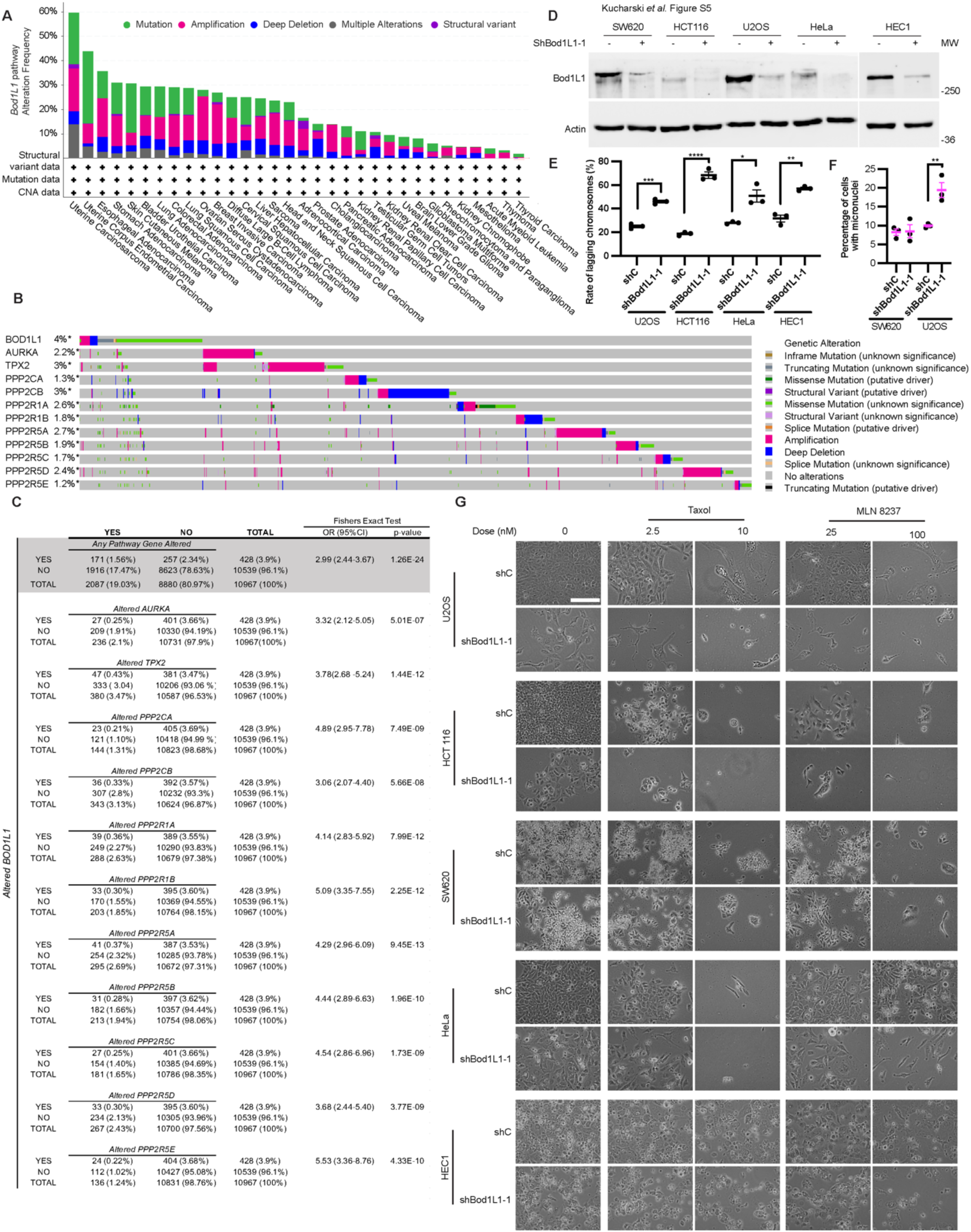
Mutations in BOD1L1 are mutually exclusive from mutations in other pathway components. A) The 10,967 sequenced cancers in the TCGA were queried for the presence of mutations in the *BOD1L1*, *Aurora A, TPX2, PPP2CA, PPP2CB, PPP2R1A, PPP2R1B, PPP2R5A, PPP2R5B, PPP2R5C, PPP2R5D, PPP2R5E* genes and plotted by cancer subtype. B) The 10,967 sequenced cancers in the TCGA were queried for the presence of mutations in *BOD1L1*, *Aurora A, TPX2, PPP2CA, PPP2CB, PPP2R1A, PPP2R1B, PPP2R5A, PPP2R5B, PPP2R5C, PPP2R5D, PPP2R5E* and plotted showing the overlap of alterations. C) Mutual exclusivity between genes in *BOD1L1* and the other queried genes was tested by *BOD1L1* alteration versus alteration to any other gene pathway member (Fisher’s exact P=1.3E-24), and *BOD1L1* alteration versus each of the eleven gene pathway members independently. D) U2OS, HCT116, SW620 and HeLa cells were plated. They were then transfected with control shRNA or shRNA against BOD1L1. 24 hours post-transfection, puromycin was added. 72 hours after addition of puromycin, the cells were harvested by scraping and lysed. The proteins were then separated by SDS-PAGE and transferred to nitrocellulose membrane and finally blotted as indicated. E) U2OS, HCT116, HEC1 and HeLa cells were plated. They were then transfected with control shRNA or shRNA against BOD1L1 and selected with puromycin. After 3 days of selection, the cells were fixed and stained with Hec1 and DAPI and the rates of lagging chromosomes were assessed from100 anaphases for each 3 independent experiments. Error bars indicate the mean +/- SEM. Statistical significance was calculated between the indicated conditions using Student’s *t-* test. F) U2OS or SW620 cells were plated. They were then transfected with control shRNA or shRNA against BOD1L1 and selected with puromycin. After 3 days of selection, the cells were fixed and stained with Hec1 and DAPI and the percentage of cells with micronuclei was determined. At least 1000 cells were examined per condition for each 3 independent experiments. Error bars indicate the mean +/- SEM. Statistical significance was calculated between the indicated conditions using Student’s *t-* test. G) Cells were prepared as in figure 7C. Following fixation, the cells were imaged by brightfield microscopy. Representative images of 3 independent experiments are shown. The scale bar is 100 μm.

**Figure S6:**
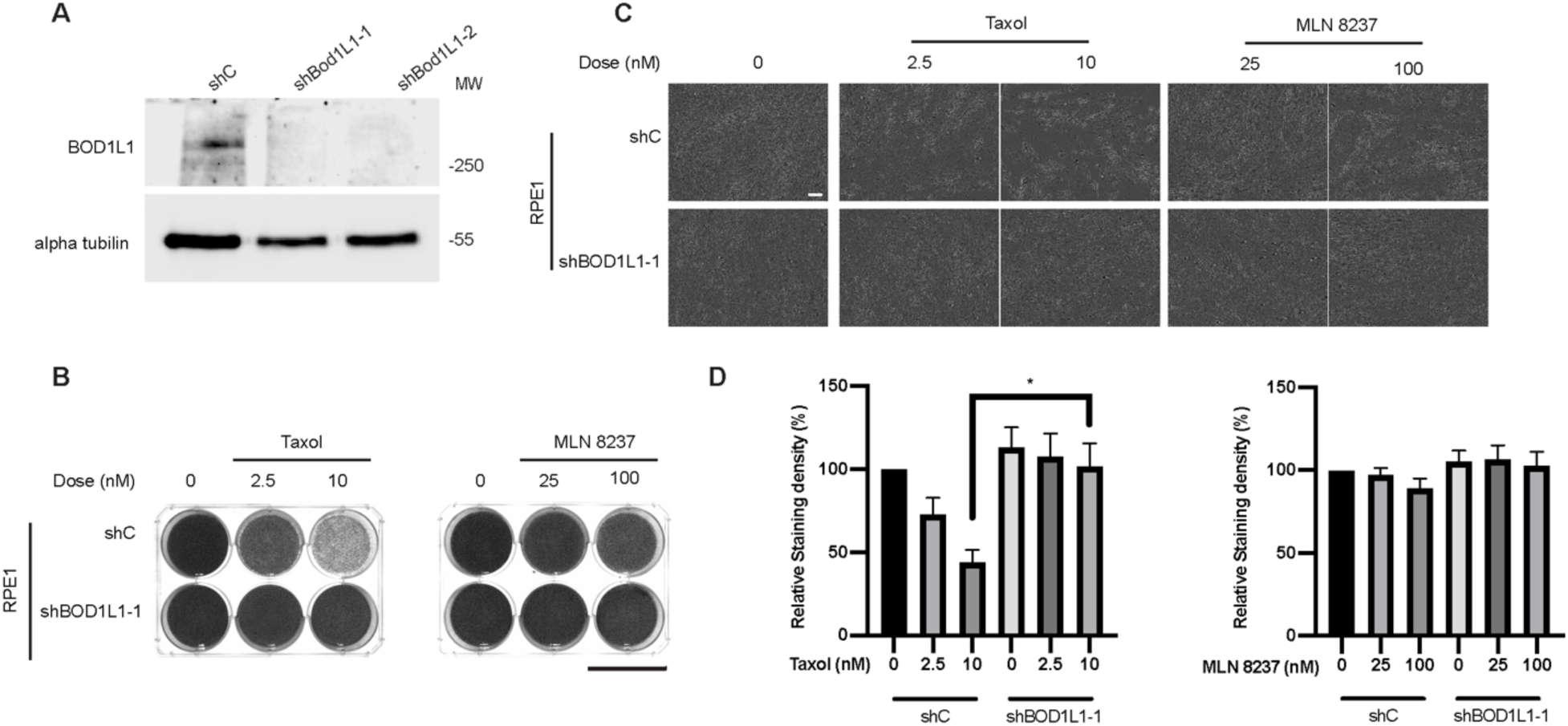
RPE1 cells are resistant to BOD1L1 knockdown. A) RPE1 cells were plated. They were then transduced with lentivirus as indicated. 24 hours following transfection, puromycin was added. The cells were harvested 72 hours after transfection and lysed. The proteins were separated by SDS-PAGE, transferred to nitrocellulose membrane and finally blotted as indicated. A single experiment was performed. B) RPE1 cells were plated They were then transduced with control shRNA or shRNA against BOD1L1 and selected with puromycin in the presence of compounds as indicated. After 1 week of selection, the cells were fixed and stained with crystal violet. Representative images from 3 independent experiments are shown. The scale bar is 4 cm. C) Cells were prepared as in figure 7C. Following fixation, the cells were imaged by brightfield microscopy. Representative images of 3 independent experiments are shown. The scale bar is 100 μm. D) Quantification of cells from (B). The staining intensity of the cells was measured using Fiji and plotted. Error bars indicate the mean +/- SEM. Statistical significance was calculated between the indicated conditions using Student’s *t-* test.

